# Osteoporosis with different causation exhibits different changes in bone tissue mineral and organic matrix

**DOI:** 10.1101/2024.12.25.630292

**Authors:** Jing-Yu Lin, Xu Guo, Ming-Hui Sun, Yifeng Zhang, Jun-Xia Lu

## Abstract

Osteoporosis (OP) which is a common skeletal disease with different causation is prevalent in aging population. Postmenopause women generally suffer from OP with bone loss due to estrogen deficiency. Diabetes are also associated with OP by complex metabolic mechanisms. Bone qualities of OP caused by aging were compared with the ovariectomy (OVX) model and the Type 2 diabetic model using Sprague-Dawley (SD) rats in our study. Combining with micro-computed tomography (*μ*-CT) and solid-state NMR (SSNMR) methods, this research studied bone changes in SD rats from tissue level to the molecular level. The studies revealed bone loss was most significant for cancellous bones, but not for cortical bones in OP rats. However, at the molecular level, the content of HAP in cortical bone increased with aging, contributing to the brittleness of the bone. Triglyceride, as a senescence maker of osteocyte in cortical bone, was also identified to be closely associated with OP in aging and OVX rats, but not in diabetic rats. This research suggests differing bone qualities in molecular level of OP with various causes more objectively reflect the bone tissue reconstruction rather than bone loss in *μ*-CT analysis.

## Introduction

Osteoporosis (OP) is a progressive disease usually associated with aging [1]. The disease affects mostly elder population, but not limited to post-menopausal women, creating a significant economic burden[2]. It is accompanied by a decrease in bone mass and an increase in bone fragility. However, bone tissue has a complex architecture with multiple components and multiscale hierarchical structures. The interplay between different components across multiscale combinatory would determine the mechanical strength and quality of the bone[1]. Therefore, more parameters reflecting changes in bone quality and quantity in OP are required to fully understand the development and different causes of OP.

Macroscopically, bone can be divided into two forms: cortical bones and cancellous bones. Cortical bone represents the strong and compact outer layer of bone tissue, while cancellous bone consists of dense trabeculae, showing spongy structures with a high surface area[3]. Bone volume, and bone density, among other parameters, have been used to monitor bone development and disease progression[4, 5]. Bone development and remodeling are dynamically controlled by the balance between the bone-forming function of osteoblasts and the bone-resorbing function of osteoclasts [6], regulating those parameters. However, clinical studies suggest that these parameters do not fully account for bone changes surpassing quantity of Ca/P compound level. Further assessment of bone at the molecular level is needed to fully characterize fine construction changes of bone.

At the molecular level, bone minerals mainly consists of hydroxyapatite (HAP) and amorphous calcium phosphate (ACP)[7, 8]. These minerals are associated with an organic matrix mainly composed of collagen (type I) fibrils[9]. These structures further assemble into microscale arrays with water canals, which are the basic structures of osteons[10]. The content of and the interaction between the inorganic and organic components also determine the quality of bone. Solid-state NMR (SSNMR) has been successfully applied in study bone and mineral materials, able to provide information on the chemical structure and dynamics quantitatively[11, 12] at the molecular level. In this paper, combined with *μ*-CT (Computed Tomography) studies, SSNMR experiments were carried out to investigate the role of aging on bone using SD (Sprague-Dawley) rats. SSNMR focused on spatial and temporal changes of minerals and organic matters in bones, comparing these changes to OP with other underlying causes.

Besides aging, there are many other risk factors associated with osteoporosis, such as estrogen deficiency[13], lipid metabolism disorders[14], diabetes[15], etc. Postmenopause women experience faster bone losses because of estrogen deficiency[13]. Ovariectomy (OVX) was used in this study to simulate postmenopausal primary osteoporosis in rats, which is a well-established model in studying OP[16]. Another animal model we studied in this paper was type 2 diabetic rat. Diabetes can have profound effects on bone health, leading to an increased risk of osteoporosis and fractures. It is reported that the higher risk is more evident for young-age individuals[17]. In type 2 diabetes, individuals can have normal to increased bone mineral density (BMD), suggesting a reduction of bone quality but not quantity, could be the reason. The abnormal bone quality may be originated from factors such as insulin resistance or hormone imbalance[18]. In our study, we carefully compared the two OP disease models with aging rats and found differences and similarities in changes in bone tissue from different aspects. For example, we observed an increase of triglyceride with OP more significantly in OVX rat than normal aging rat, but not in diabetic rat. OVX rat also has extremely low content of HAP and ACP, probably caused by the immature development of bones. Upon aging, the cancellous bone shows a significant decrease in bone volume and density from 10 months of age, causing OP. However, our NMR studies show that the mineral HAP continues to evolve and increase until 21 months of age, increasing the crystallinity of bone, which is more significant for cortical bone and skull than cancellous bone. For the diabetic model, a consistent decrease of ACP content is observed for all three types of bones, also leading to the increase in HAP contents in bone, causing the increased crystallinity and brittleness of bone. The information gained in this study provides insight to help us understand OP with different underlying mechanisms, which will provide us new means to the early detection of OP before a fracture occurs and is essential in developing medicine treating OP.

### Experimental procedures

#### Sample preparation

##### Animals

All animal procedures were carried out in accordance with the Ethics Committee and approved by the Administrative Panel on Laboratory Animal Care (protocol no. 20210914001) of ShanghaiTech University. Samples were collected on the femur (mid-diaphysis), lumbar (third lumbar vertebra) and skull (bilateral parietal bone) of SD (Sprague-Dawley) rats with different ages (1/2/6/10/21/32 months). All samples studied in this research are shown in figure 1.

**Figure 1.**
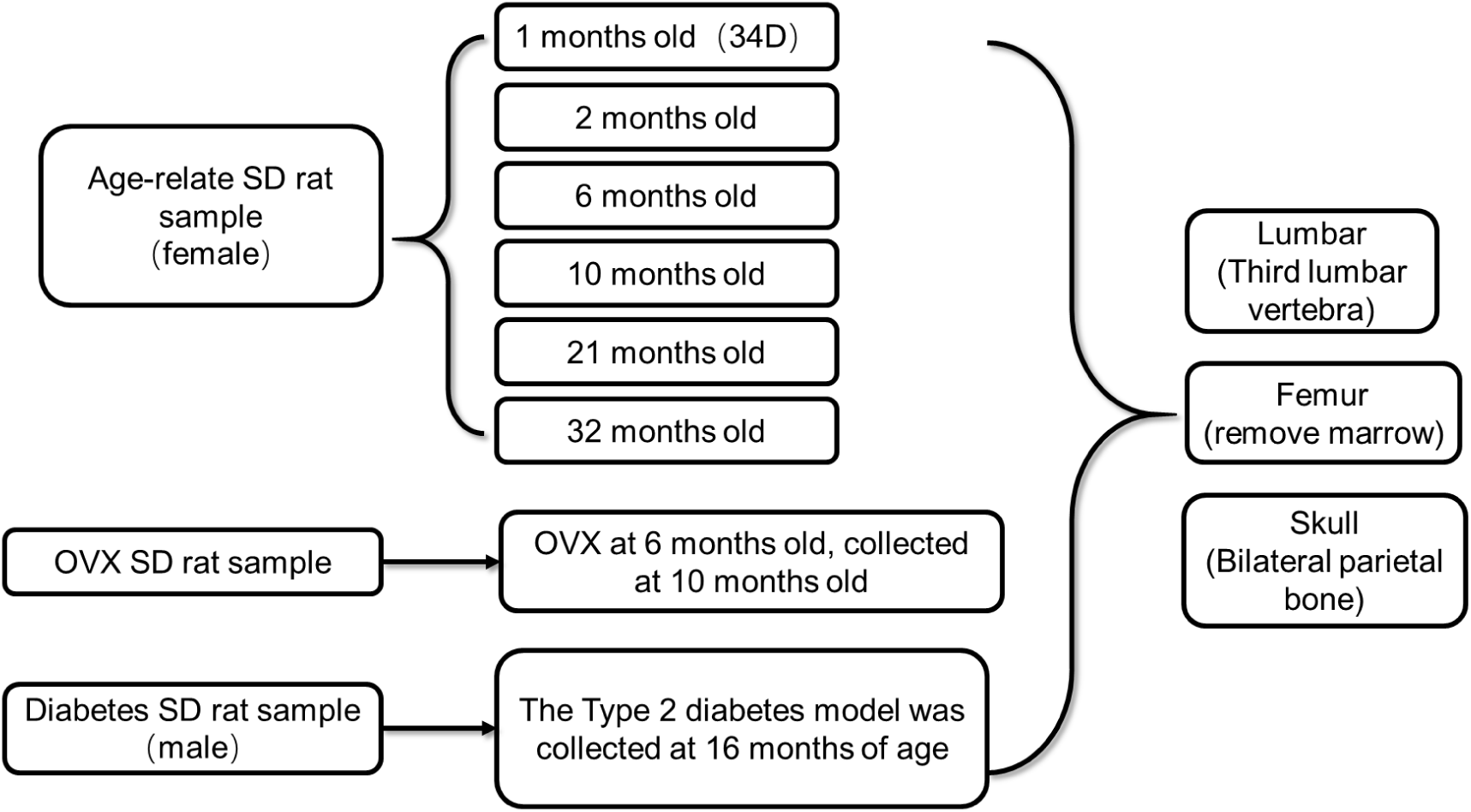
Bone tissue samples studied in this research. SD rats aged from 1 to 32 months were used in age-related studies. An OVX rat and type 2 DM rat bone samples were also investigated to study OP induced by different causes. The bone tissues studied include the third lumbar vertebra, the mid-diaphysis femur, and the bilateral parietal bone of the skull. All samples were analyzed by *μ*-CT and SSNMR.

##### Ovariectomy rat model

The female rats were operated on at 6 months of age to remove ovarian tissue and continued to be cultured until 10 months of age for *μ*-CT and SSNMR detection. First, the lower margin of the ribs was incised with a 0.5cm-1cm opening, located 1cm away from the spine. Following the dissection of the psoas muscle, ligation was performed on the fallopian tubes at the upper portion of the uterine horn and the ovarian tissue was removed. The cellulite was repositioned, the muscle and skin were sutured, and penicillin was administered via injection for sterilization. The other ovary underwent a similar removal procedure. Once awake, the rats were returned to their cages.

##### Type 2 diabetes SD rat model

A type 2 diabetes rat model was established by inducing type 2 diabetes mellitus (DM) in male SD rats aged 6 weeks through intraperitoneal injection of a low dose of streptozotocin (STZ) and feeding them a high-fat diet (China Jiangsu Xitong Pharmaceutical Bioengineering Co., LTD, consisting of 23.7% protein, 41.4% carbohydrate, and 23.6% fat). After being on the high-fat diet for 4 weeks, the rats were injected with streptozotocin (STZ (30 mg/kg)) via intraperitoneal route. Weekly measurements of blood glucose levels were taken in DM rats, and STZ (30 mg/kg) was administered again to those with glucose concentrations lower than 16.7mmol/L. Four weeks after the initial STZ injection, rats with blood sugar levels >16.7 mmol/L were considered diabetic and continued on a high-fat diet until they reached the age of 16 months[19].

### *Micro*-Computed Tomography (*μ*-CT)

The femur, skull and lumbar were separated and scanned by SkyScan1276(Bruker). Scans were conducted with a source voltage of 70 kV and a source current of 200 μA. Scanning parameters were set as follows: Filter: Al 1.00mm; Binning: 2016×1344; pixel size: 20.36951 μm; X-ray radiation source to bone distance: 158.548 mm. The scan was reconstructed in the range 0.006417-0.069248. The data were analyzed with instrumental software CTAn. In the analysis of lumbar, the area of the third lumbar vertebra was selected for analysis; In the analysis of the skull, the skull was placed vertically, according to the height of the sagittal suture as the analysis range, half of the sagittal suture was taken as the side length of the area of interest, and the width of the area of interest was determined according to the thickness of the skull. In the analysis of the femur, 100 slices were selected above and below the mid-diaphysis, totaling 200 slices. The size of each cross-sectional area in the region of interest was 200 mm×200 mm. Within these defined regions of interest, bone morphometric parameters of percentage bone volume/tissue volume (BV/TV, %), bone mineral density(BMD) and BMD mean were measured. BMD mean is different from BMD in that BMD mean is calculated on the bone density of hard tissue only, calculated by removing the soft tissue volume within the region of interest. BMD is calculated on the whole bone tissue.

### SSNMR sample preparations

All the bone samples (femur, lumbar and skull) were collected immediately after *μ*-CT scan and freeze-dried. Femoral samples were ground into powder after removing the marrow. Lumbar samples were further removed from the muscle tissue after draining and later ground into powder. Skull samples were ground into powder directly. Each sample was accurately weighed. Type I collagen from bovine achilles tendon (Collagen Type I; Solarbio; China) was also freeze-dried and ground for SSNMR analysis.

### SSNMR experiments

All the SSNMR experiments were acquired on a 16.4-T (700 MHz ^1^H frequency) Bruker AVANCE NEO spectrometer with 3.2-mm triple-resonance HCP or HCN magic angle spinning (MAS) probe, and the MAS speed was 15kHz. All experiments were performed at 288K for HCP probe and 293K for HCN probe. ^1^H chemical shift was indirectly referenced to sodium trimethylsilylpropanesulfonate (DSS) at 0 part per million (ppm) and ^13^C chemical shift was externally referenced to DSS by setting a downfield ^13^C signal of adamantine to 40.48 ppm[20]. For HCP probe, the ^31^P chemical shifts were indirectly referenced to 85% H_3_PO_4_ (0 ppm) using the ^31^P/^1^H resonance frequency ratio 0.404807420. All of the experiments were collected with a 2s recycle delay.

For experiments on HCN probe, ^1^H 90-degree pulse was 3.35μs with the radiofrequency (rf) strength of 74.63 kHz, and ^13^C 90-degree pulse was 3.75μs with the rf strength of 66.67 kHz. For ^1^H-^13^C cross-polarization (CP) MAS spectrum, the ^13^C rf strength was set to 44.35 kHz, and ^1^H rf strength was set to 86.52 kHz with 200μs contact time. ^1^H decoupling of 74.63 kHz was applied using spinal-64[21]. The ^1^H–^13^C frequency-switched Lee-Goldberg heteronuclear correlation (FSLG-HETCOR) [22, 23]experiment was carried out using 100μs contact time. The Insensitive Nuclei Enhanced by Polarization Transfer (INEPT) [24]spectra was recorded using ^1^J_CH_ 145 Hz and waltz16_18 ^1^H decoupling of 5.0 kHz [25].

For experiments on HCP probe, the ^1^H 90-degree pulse was 3.10μs with the rf strength of 80.65 kHz. The ^13^C 90-degree pulse was 3.00μs with rf strength of 83.33 kHz. The ^31^P 90-degree pulse was 3.13μs with the rf strength of 79.81 kHz. For 1D ^1^H-^13^C CP MAS[26], the ^13^C rf strength was set to 50.9 kHz, ^1^H rf strength was set to 79.81 kHz. The CP contact time was 200μs. For ^1^H-^31^P CP MAS, the ^31^P rf strength was set to 45.43 kHz and ^1^H rf strength was set to 78.55 kHz with a contact time of 1.5 ms. ^1^H SPINAL-64 (individual pulse width = 6.2μs) decoupling was applied during the evolution and acquisition periods. For two-dimensional (2D) ^1^H-^31^P heteronuclear correlation (HETCOR), SSNMR spectra were acquired using a ^1^H π/2 pulse of 3.28 μs corresponding to a rf of 77.9 kHz [27] with a contact time of 1.6ms. The CP match was set with a 70 %–100 % ramp-shaped pulse of 75.4 kHz on the ^1^H channel and a square-shaped pulse of 52.3 kHz on the ^31^P channel. The number of scans of SSNMR experiments and the mass of each sample are shown in Table S1 and Table S2.

Car-Purcell Meiboom-Gill(CPMG) echo pulse was applied for the ^1^H transverse relaxation time measurements(T_2_) with ^1^H π/2 pulse of 3.35μs and π pulse of 6.7μs and 12s relaxation delay. The inter-pulse delays were varied: 4ms, 8ms, 16ms, 32ms, 128ms, 256ms, 512ms and 1024ms. The signal decay was fit by the single exponential function with reasonable confidence using Origin (OriginLab Corporation, Northampton, MA USA).

### The analysis of mineral HAP and ACP content change

The ^1^H slice of the 2D HETCOR was taken and integrated from-2 ppm to 20 ppm. MestReNova program (Mestrelab Research, S.L., Spain) was used to deconvolute the slice into three components, with OH/PO_4_^3-^ (HAP) near 0 ppm, H_2_O/PO_4_^3-^ (ACP) near 5.8 ppm, and HPO_4_^2-^ near 11 ppm. NMR peak integral values (A) for these three components were obtained. The integral value per unit mass was further calculated (A/ M, (A) divided by each NMR sample mass (M)). Then it was combined with BMD mean to obtain the integral of the NMR signal per unit of bone volume for HAP/ACP/ HPO_4_^3-^ (I=BMD mean A/M) (Table S3). (BMD mean was obtained from *μ*-CT experiments and indicated the bone density of the hard tissue by removal of the soft tissue in the calculation. Although BMD mean value could be different upon bone grounding for NMR studies, it was assumed that the sample grounding process would induce similar changes for the same type of bones (lumbar, femur, or skull). Therefore the (I) values would still consistently reflect the amount of mineral changes within the same type of bone tissues.) The (I) values were used as parameters to show the changes of the three mineral components with OP development. In addition, the proportion of HAP to the total amount of HAP and ACP was calculated from the peak integral ratio between the two components.

## Results

### Aged-related changes in cortical and cancellous bone revealed by *μ-*CT

Femur (mid-diaphysis), lumbar (third lumbar vertebra) and skull (bilateral parietal bone) were taken from SD rats with different ages (1/2/6/10/21/32 months) and a comparative analysis was conducted using *μ*-CT. The mid-diaphysis of femur was analyzed as the representation of cortical bone (Fig. 2) while the third lumbar vertebra was analyzed as a representation of cancellous bone (Fig. 3). The skull, representing a combination characteristic of both cortical and cancellous bones, was analyzed against the sagittal suture (Fig. S1).

**Figure 2.**
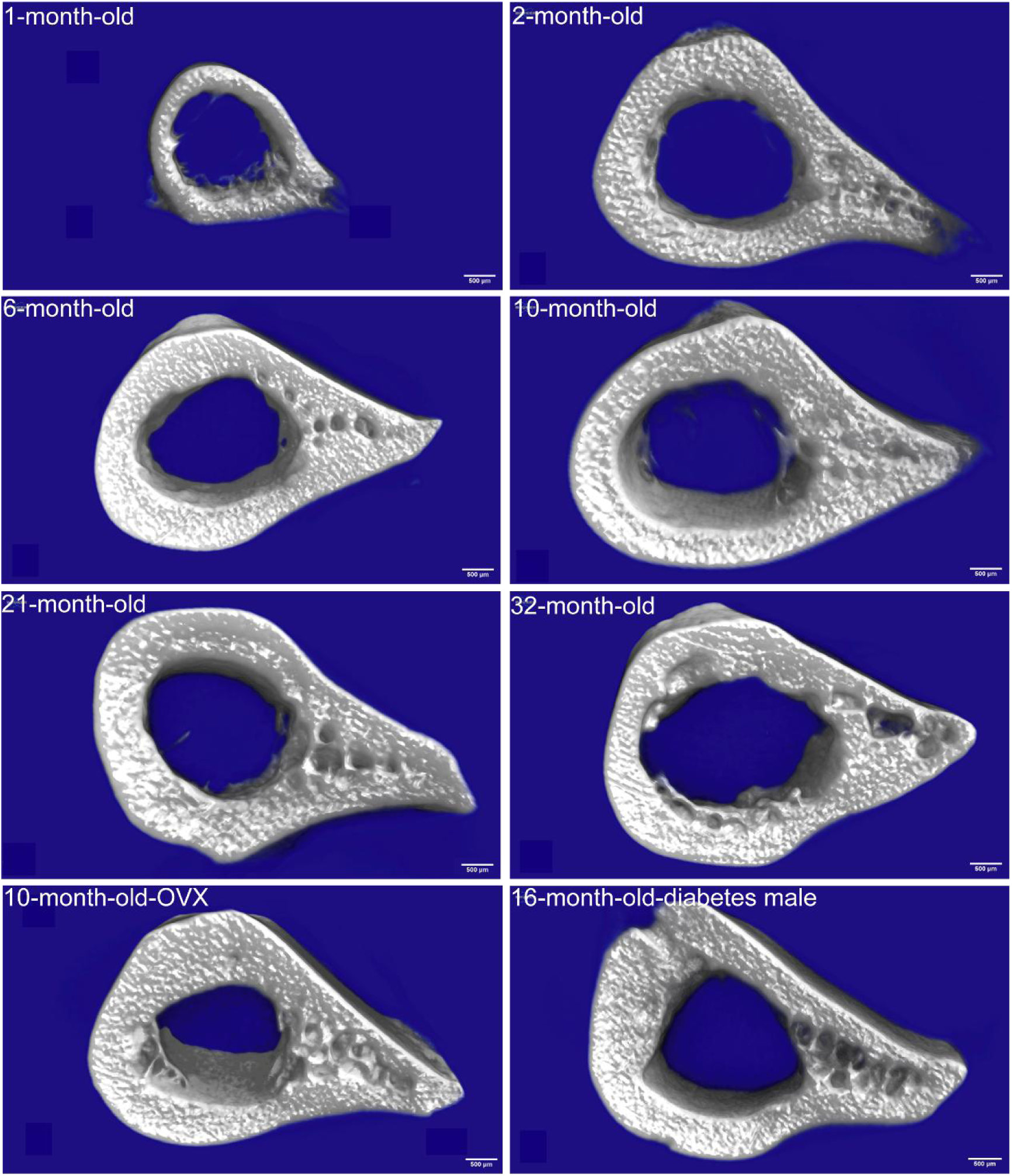
The *μ*-CT images of the cortical bones represented by femur, showing a progressive change of bone with age and OP. The sample was from 1, 2, 3, 6, 10, 21 and 32-month-old SD rats, the10-month-old-OVX rat and the 16-month-old male SD rat with diabetes. The scale bar in each image is 500μm.

**Figure 3.**
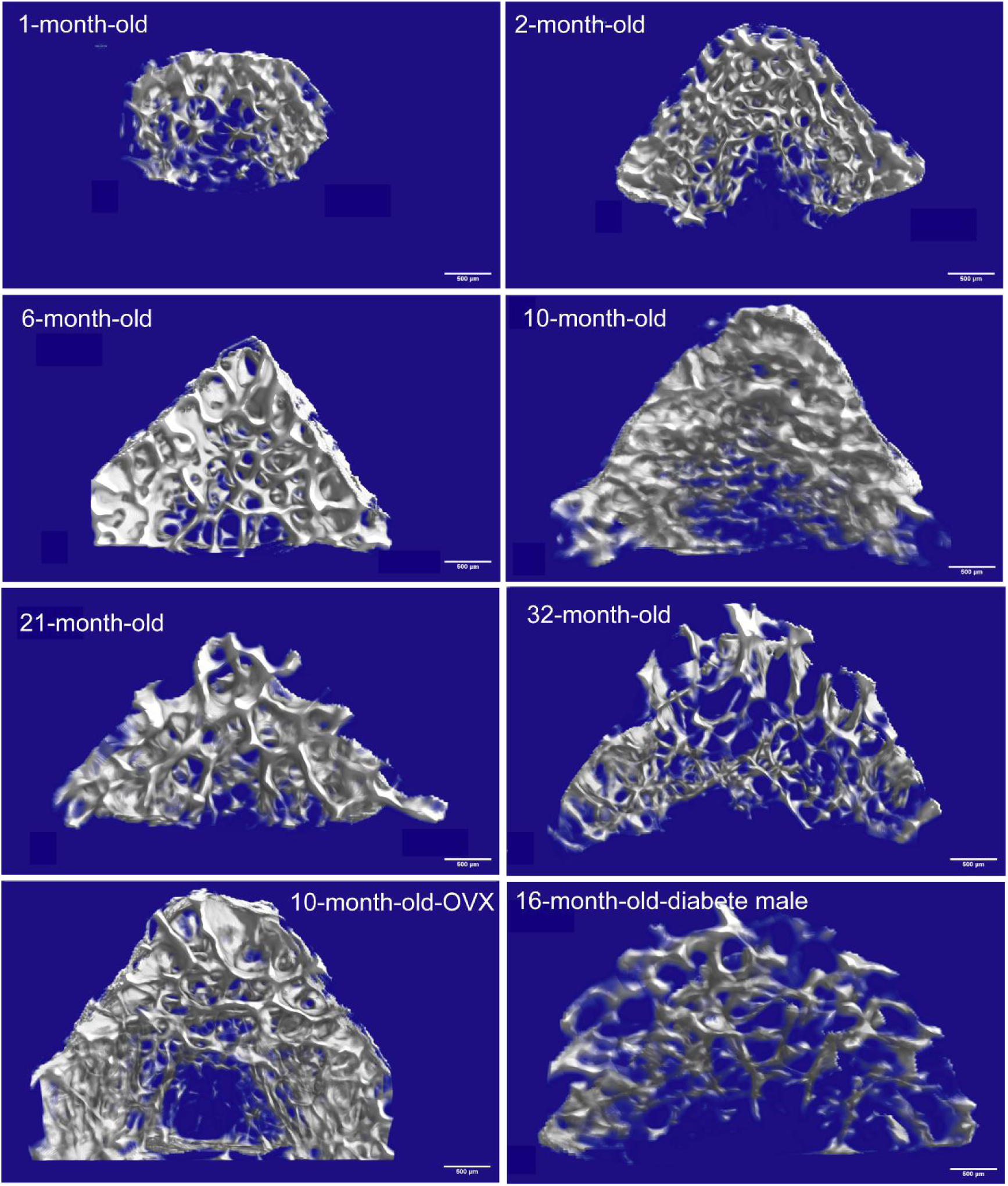
The *μ*-CT images of the cancellous bones represented by lumbar, showing a progressive change of bone with age and OP. The sample was from 1, 2, 3, 6, 10, 21 and 32-month-old SD rats, the 10-month-old-OVX rat and the 16-month-old male SD rat with diabetes. The scale bar in each image is 500μm.

The transverse cross-sectional image of rat femoral diaphysis shows a gradual increase in the thickness of cortical bone and an enlargement of marrow cavity up to about the age of 6-10 month (Fig. 2). As the rats aged, bigger pores indicative of osteoporosis emerge in cortical bone. *μ*-CT reveals low BV/TV (bone volume/tissue volume) and BMD (bone mineral density) during early cortical bone growth at 1 month, followed by significant elevation of both parameters due to increased osteogenic activity from 1 to 6 months. However, changes plateau after 6 months, with BV/TV and BMD declining only slightly during 10 to 32 months (Fig. 4A, B).

**Figure 4.**
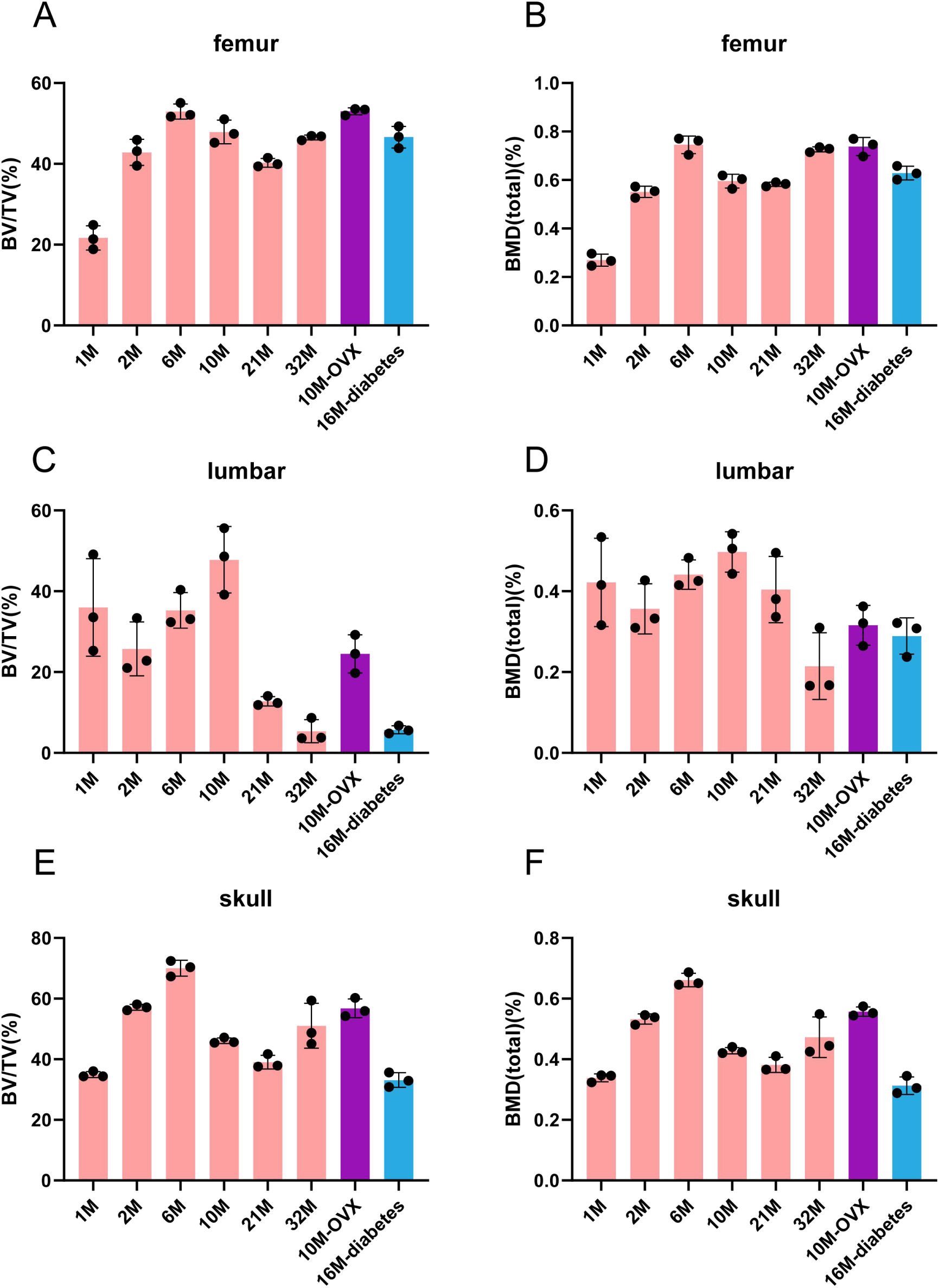
The morphometric analysis of three types of bone tissues by *μ*-CT. The change of bone volume/tissue volume (BV/TV) is shown on the left for (A) femur, (C) lumbar and (E) skull. The change of bone mineral density in total volume (BMD (total) (%)) is shown on the right for (B) femur, (D) lumbar and (F) skull.

Concurrently, the third lumbar vertebra representing cancellous bone shows a gradual increase in size indicating bone growth (Fig. 3, 1-10 months). However, larger voids and thinner trabecular bone in old age suggest the development of osteoporosis (Fig. 3, 32 months). Quantitative analysis by *μ*-CT reveals BV/TV and BMD peaked at 10 months, followed by a clear decline, with evident osteoporosis at 32 months (Fig. 4C, D). Therefore, cancellous bone, being porous and stress-dispersing, is more susceptible to bone loss due to aging[28–30].

The skull contains both a cortical and cancellous bone portion. At 1 month of age, the skull is not yet fully developed. The *μ*-CT findings indicate a large gap in the sagittal suture of the bone (Fig. S1). The cranium undergoes rapid ossification and is structurally more complete between 2 and 6 months of age (Fig. S1). Thereafter, small pores in the skull gradually increase (Fig. S1, 10-32 months). Quantitative analysis using *μ*-CT shows BV/TV and BMD changes similar to femur, but different from lumbar (Fig. 4E, F). At 1 month of age, BV/TV and BMD of skull are at relatively low levels. When the rats grow to 6 months of age, both BV/TV and BMD reach their peaks. After 6 months of age, BV/TV and BMD begin to decline but to a less extend compared to cancellous lumbar bone (Fig. 4E, F).

### Aged-related changes of minerals in cortical and cancellous bone revealed by SSNMR

NMR sample preparation was performed on the samples immediately after the *μ*-CT experiments. Bone marrow and muscle tissue were removed from bone before it was ground into powder. All samples were freeze-dried to remove the free water. 2D ^1^H-^31^P HETCOR experiments were carried out on the three types of bone samples (Fig 5, Fig S2, S3), showing the changes of bone as SD rats aged. The 2D ^1^H-^31^P HETCOR experiment results show the presence of three different localized phosphate environments in correlation with ^1^H: the correlation peak between PO_4_^3-^ and OH^-^ ions (^31^P/^1^H ∼3.6/0.5 ppm) indicating the presence of hydroxyapatite crystals (HAP), the correlation peak of phosphate groups with water molecules (^31^P/^1^H ∼3.6/5.9 ppm) which represents the amorphous calcium phosphate (ACP) association with trace tightly bound water on mineral surfaces, and the peak associated with HPO_4_^2-^ or H_2_PO_4_^-^ (^31^P/^1^H ∼3.6/12 ppm)(Fig 5, Fig S2, S3). A deconvolution of ^1^H slice of 2D ^1^H-^31^P HETCOR was shown in fig 6A as an example, indicating the three phosphate mineral components. 1D slices were also displayed on the 2D spectra, and compared to their corresponding 1D ^1^H direct pulse MAS spectra, indicating a significant difference. 1D ^1^H MAS spectra usually exhibited peaks centered around 1-2 ppm (from protein or lipids acyl chain) and a broad peak at 5.3 ppm (mainly from associated water not removed by lyophilization).

**Figure 5.**
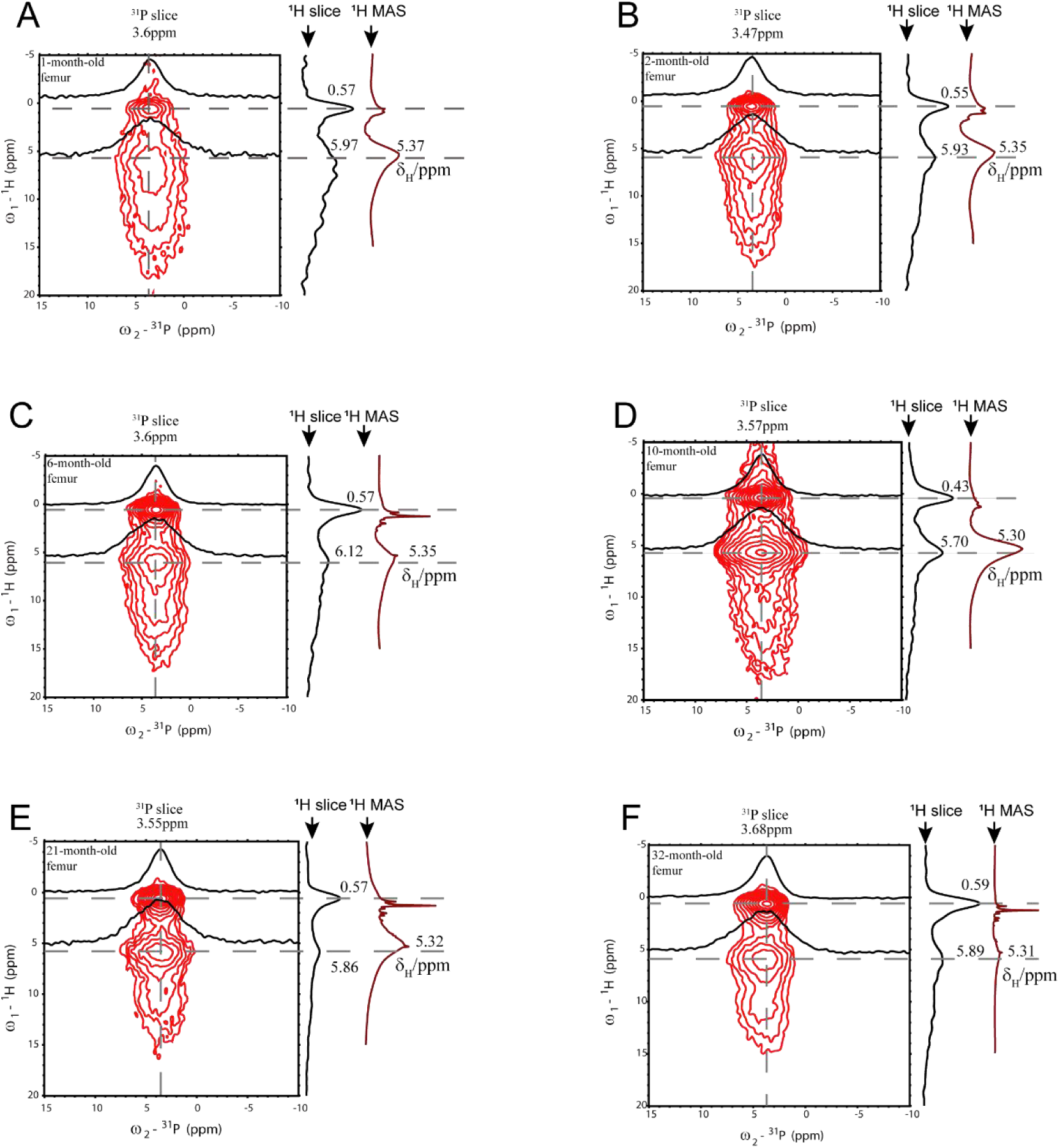
2D ^1^H-^31^P HETCOR spectra of the cortical femur bones. A, B, C, D, E and F represent femur from 1, 2, 3, 6, 10, 21 and 32-month-old SD rat, respectively. All femur samples were freeze-dried, and the spectra were collected from 128 scans.

The deconvolution in fig 6A allows us to quantitatively analyze the differences of these 2D spectra in the content of these three mineral components (Table S3, Fig 6B, 6C, 6D). Quantitative analysis showed that in cortical bone (femur), the content of HAP (OH^-^/PO_4_^3-^) increases by about 415% between 1 and 21 months of age, then enters a plateau period and only decreases slightly at 32 months of age. ACP increases by about 86% between 1 and 10 months of ages but stabilizes after 10 months of age and reaches a plateau (Fig. 6B) without significant decay. This observation on ACP is consistent with the observed trend on μ-CT results, which show only a small decrease in BV/TV and BMD from 10 months of age for cortical bone. But the change trend of HAP in cortical bone reaching peak at 21 months of age is different from the observed trend on BV/TV and BMD. The linewidths of the ^31^P CP spectra under conditions of high ^1^H power decoupling correlate with the localized heterogeneity of the chemical environment in Ca/P minerals[11]. Analysis of the 1D ^31^P CP spectra (Fig S4, Table S4.) reveal an overall decreasing trend in the linewidth of the cortical bone (Fig. 7A). And the proportion of HAP in total HAP and ACP intensity, calculated from fig 5, increases by 65% between 1 and 32 months of age (Fig. 7B). As HAP represents a much ordered structure than ACP, these results combined support an increase in the crystallinity of cortical bone mineral upon aging.

**Figure 6.**
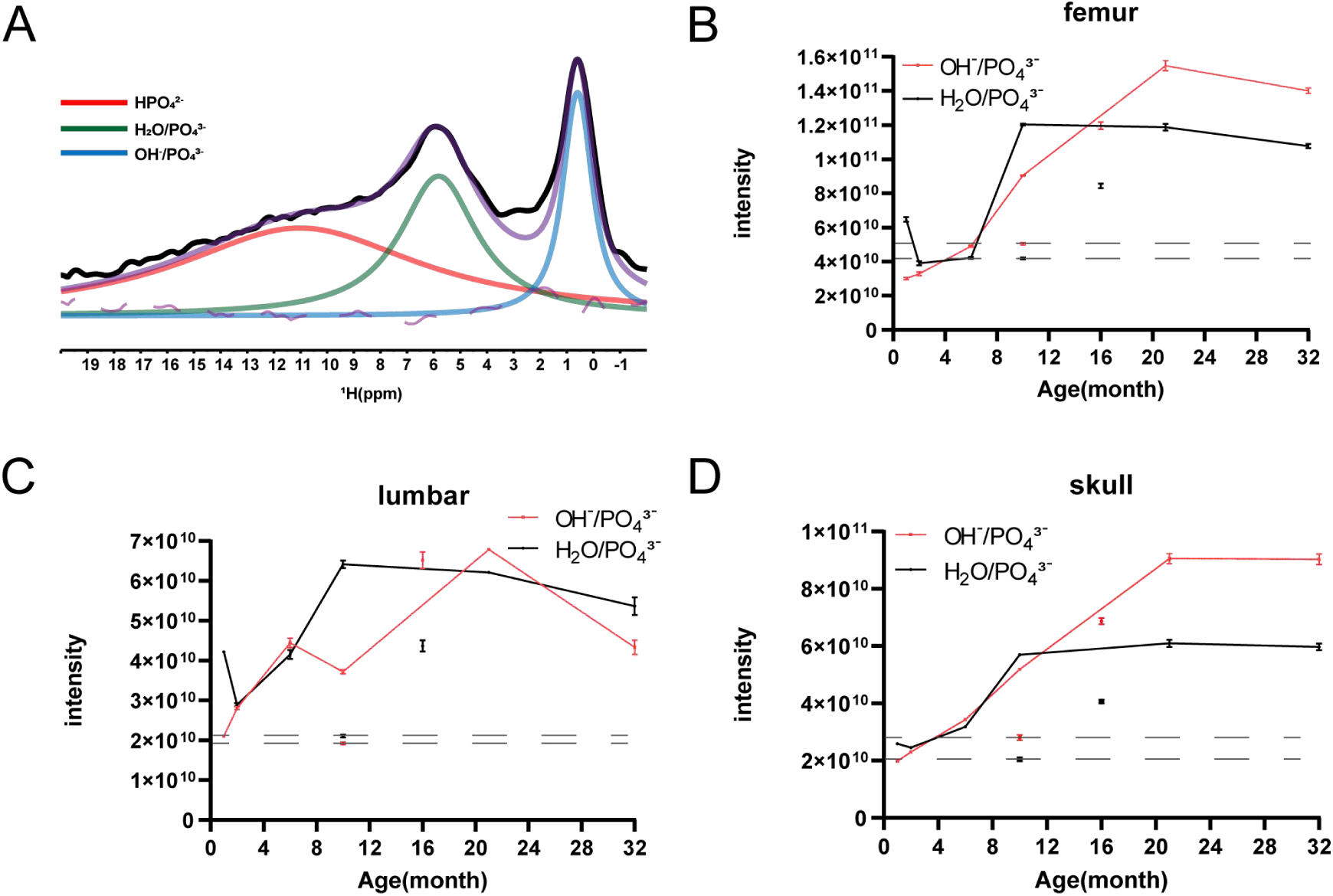
The changes of HAP and ACP were detected from bone samples by a combination analysis of both SSNMR and *μ*-CT results. **The intensity of** different components of H 1D slices from H- P 2D HETCOR was used to obtain the content change information. (A) The H 1D slice from 2D HETCOR of 10-month-old lumbar was shown as an example here. Red: HPO_4_ ^2-^; Green: H O/PO ^3-^; Blue: OH/PO_4_ ^3-^; Dashed line: error between fitted data and experimental data; Black: experimental data; Purple: Fit curve. (B) Integral intensity of H O/PO ^3-^ and OH/PO_4_ ^3-^ per unit bone volume changes as rat aging or OP development for femur, (C) lumbar and (D)skull. For the disease model, the results for the 10-month-old-OVX rat samples and the 16-month-old male SD diabetic rat samples were also shown as points at the corresponding ages in the figure. The data are shown in Table S3. The values for the 10-month-old-OVX rat samples were also marked with dashed line.

**Figure 7.**
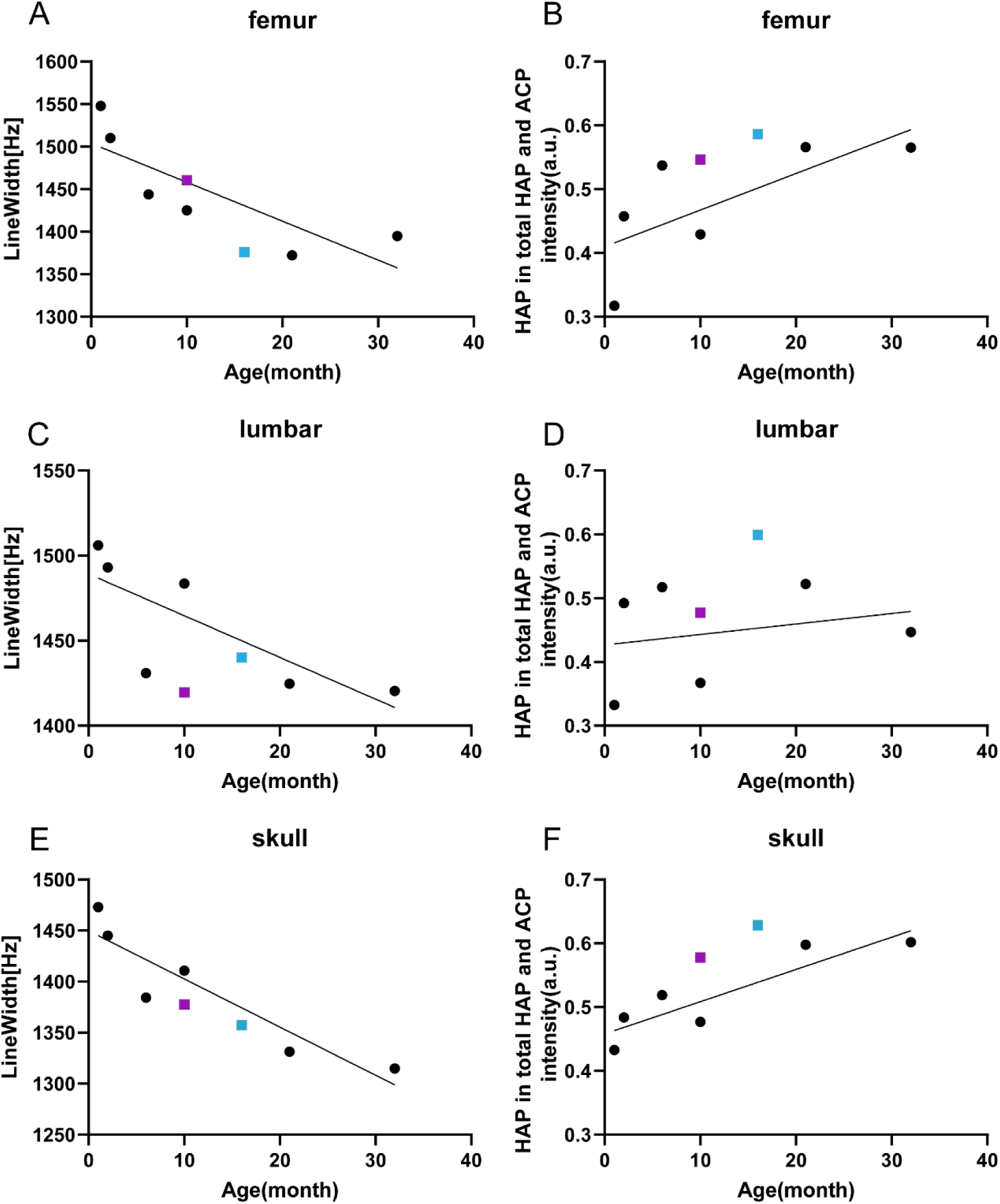
The ordered CaP structures in cortical bone increased with age. The changes of ^1^H-^31^P CP linewidth for (A) femur; (C) lumbar (E) skull samples. The increase of HAP estimated by the percentage of HAP in total HAP and ACP for (B) femur; (D) lumbar (F) skull. The results from two disease models were shown at the corresponding age. Purple: the 10-month-old-OVX rat; Blue: the 16-month-old male diabetic SD rat.

In cancellous bone (lumbar), the overall trend of HAP is like that of cortical bone. The amount of HAP increases generally until the rat growing to 21 months of age but decreases slightly between 21 and 32 months (Fig 6C). Similarly, as the cortical bone, ACP increases until the rat growing to 10 months of ages but decreases by ∼16% between 21 and 32 months (Fig. 6C). However, the intensity of HAP and ACP keeps at the similar levels. There is also an overall decreasing trend in the linewidth of cancellous bone in the 1D ^31^P CP spectrum (Fig. 7C). At the same time, the proportion of HAP in total HAP and ACP remains similar or only slightly increases between 1 and 32 months of age, judged by the slope in fig. 7D. Therefore, the overall increase in the bone mineral crystallinity is much smaller than the cortical bones.

2D ^1^H-^31^P HETCOR spectra were also obtained on the skull samples (Fig. S3). Quantitative analysis show that HAP has an increasing trend from 1-21 months of age, increasing by approximately 358%, and reaches a plateau after 21 months of age (Fig. 6D). In contrast, ACP shows an overall increasing trend from 1-10 months of age, with an increase in content of ∼120%, and reaches a plateau at 10 months of age (Fig. 6D). The results are consistent with those obtained from femur samples. In the 1D ^31^P CP spectra, there is an overall decreasing trend in linewidth (Fig. 7E), and the proportion of HAP to total HAP and ACP increases by 40% from 1-32 months of age (Fig. 7F). Therefore, the overall increase in the bone mineral crystallinity for cortical bones and skull (with a mixture of both cortical and cancellous bone properties) upon aging can be concluded, while it is less obvious for cancellous bones.

Considering cancellous bone (lumbar) μ-CT results were very different from cortical bone (femur) and skull for older SD rats, showing a significant decrease in BV/TV and a clear decline in BMD after 10 months, SSNMR revealed information at the molecular level and exhibited similar changes of HAP and ACP for femur and skull, but different change for lumbar (Fig. 6). In lumbar, the intensities of HAP and ACP are similar. But in other two bones, the increase of HAP continues at older age (21 month) while ACP plateaus at 10 months, leading to a higher HAP content than ACP at older age. SSNMR also exhibits that the mineral intensity per unit bone volume is much higher for femur bone than skull and lumbar, with lumbar bone containing the least minerals by comparing the intensity scale in fig 6B, C, D. The results are also consistent with the *μ*-CT results, showing the cortical femur bones are denser in mass and contain more minerals than cancellous bone.

### Aged-related changes of the organic matters in cortical and cancellous bone revealed by SSNMR

The changes on the organic matrices reflect a change of the mineral microenvironment, giving clues on bone mineral development. To understand the changes of organic matrices during rat aging, we conducted 1D ^1^H-^13^C cross-polarization (CP) experiments with a focus on the abundant type I collagen within bone tissue. All spectra were compared to the 1D ^1^H-^13^C CP spectrum of purchased type I collagen from bovine achilles tendon. While signals of collagen are evident across all bone tissue samples, no notable distinctions emerge in the profiles of cortical and cancellous bone with different ages (Fig. 8A, B). Following quantitative analysis through integral processing of the spectra, we observed a decrease of approximately 15% intensity within cortical bones (Fig. 8C) and an increase of approximately 50% within cancellous bone between 1 and 32 months of age (Fig. 8D). For skull, the content of type I collagen decreases between 1 and 32 months of age (Fig. S5A, B), following a similar trend as in cortical bone (Fig. 8C). Therefore, type I collagen content in bone is affected by age and differs cumulatively in cortical versus cancellous bone. An increase in the spectra intensity of aging cancellous bone could probably be caused by the significant decreased extracellular Ca/P deposition with decrease of bone volume and density, leaving extra organic matrix unsaturatedly mineralized.

**Figure 8.**
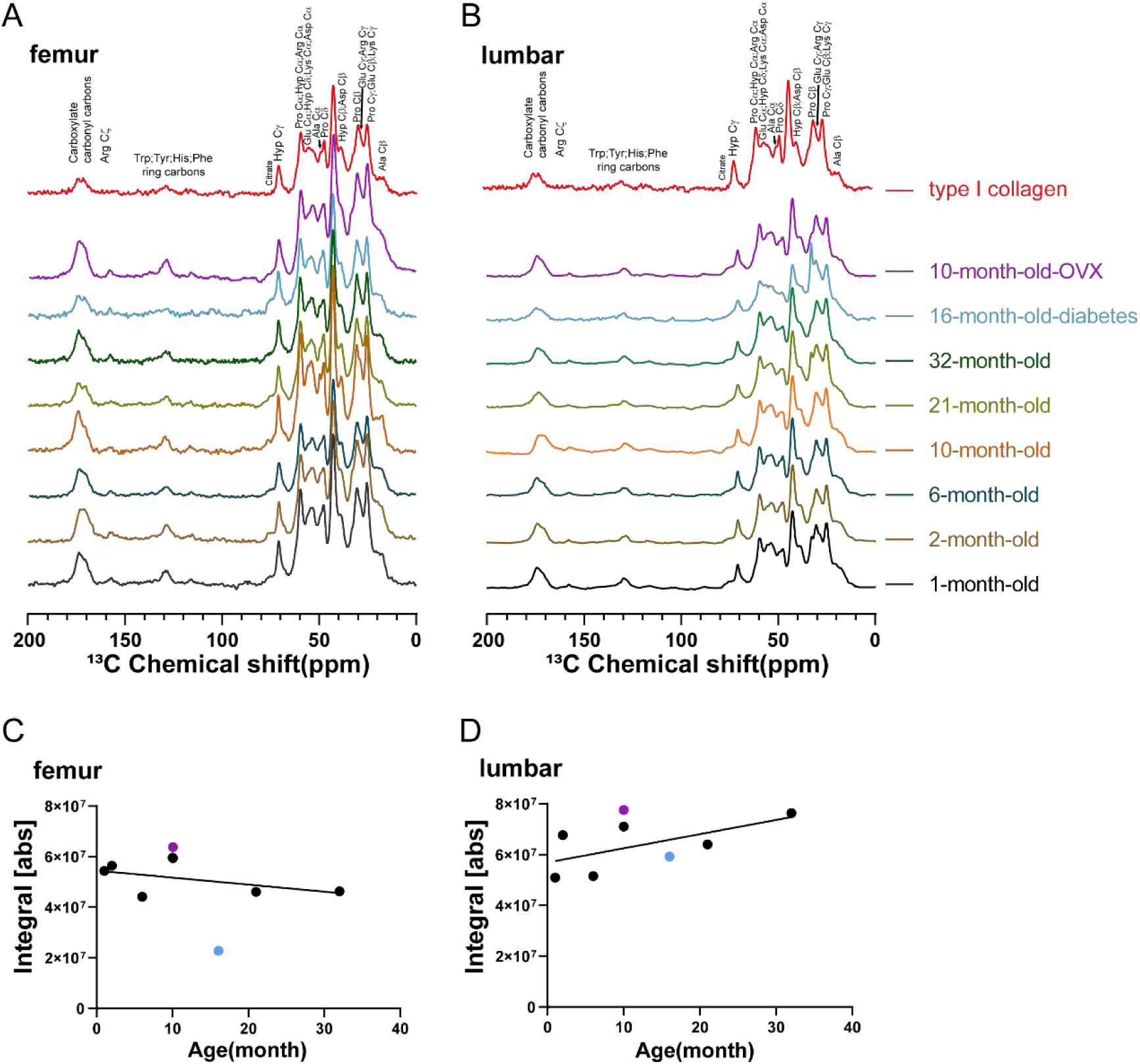
Comparison of 1D ^1^H-^13^C CP SSNMR spectra of SD rat bone samples, exhibiting a typical spectrum of type I collagen. (A) is for femur and (B) is for lumbar samples with different ages. The 1D ^1^H-^13^C CP spectra measured at 15kHz MAS. In (C) femur and (D) lumbar, the spectral integrals were shown for bones with different ages. All the spectral integrals are normalized by sample mass and the integral range is 0 ppm∼200 ppm. The results on two disease models were also shown. Purple: the 10-month-old-OVX rat; Blue: the 16-month-old male diabetic SD rat.

At the same time, we conducted a comprehensive examination of all hydrogen-containing components using the 1D ^1^H DP experiment. The results show that the spectral signals at 0-3 ppm were weak at 1 ∼ 10 months of age and are significantly stronger at 21 ∼ 32 months of age, especially for cancellous lumbar bone (Fig. 9A, B). Interestingly, there is no such an increasing trend of ^1^H DP intensity at the same spectral range for skull while rats aged (Fig. S5C, S5D). The major peaks for ^1^H DP spectra are at 0.88, 1.29, 1.56, 2.04, 2.24, 2.79, and 5.34 ppm. These signals overlap with some ^1^H signals of type I collagen and are also consistent with those of other molecules, such as triglycerides and lipids, and thus could not be accurately determined. Considering 1D ^1^H-^13^C CP signals of collagen in bone tissue are not changed significantly, these signals are likely not from rigid collagen fibrils in the bone organic matrix.

**Figure 9.**
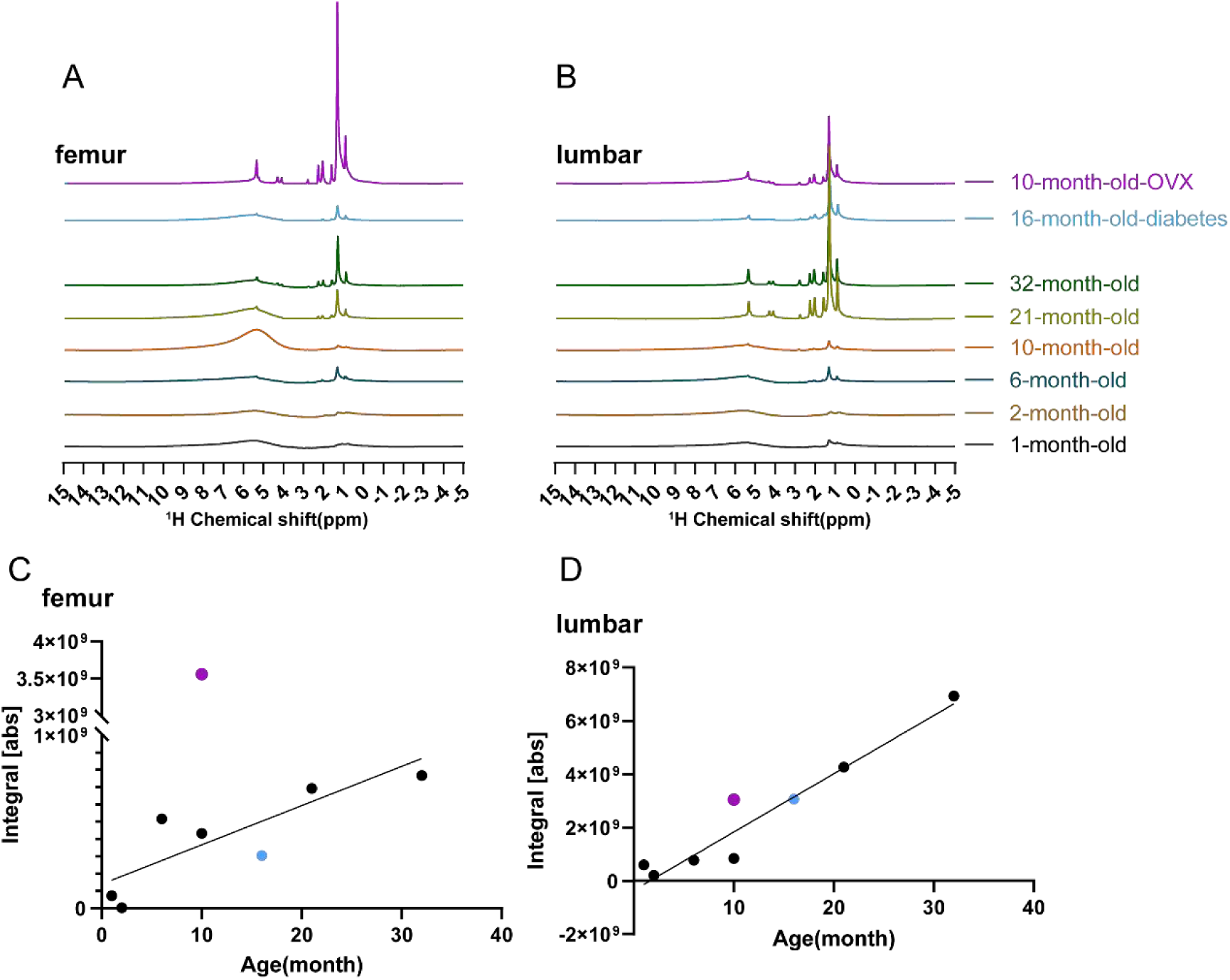
1D ^1^H DP spectral changes with age and diseases. 1D ^1^H DP spectra for (A) femur and (B) lumbar. All the spectral intensities were normalized by mass. The peak integral (0∼3 ppm) changes were shown in (C) femur; (D) lumbar. Purple: 10-month-old-OVX SD rat; Blue: 16-month-old male diabetic SD rat.

To analyze the more mobile organic matter in bone, 1D ^1^H-^13^C INEPT experiment was carried out. The results show that the changes in INEPT signals in bone upon aging are consistent with the 1D ^1^H DP results, indicating the increase of signals of DP spectra at older age are originated from the more mobile organic substances (Fig. 10). T_2_ values were also obtained on ^1^H DP spectrum for the 32-month-old lumbar sample, showing long T_2_ values from 15-30 ms, consistent with a high mobility of the molecule (Table S5, Fig. S6). Furthermore, although these substances are prevalent in both cortical and cancellous bone, the signal intensity is greater in cancellous bone. Again, no such increase was observed for skull. These results indicate that the three types of bone show different changes regarding this mobile organic component. Finally, 2D ^1^H-^13^C INEPT was used to identify the mobile organic component. By comparing the spectra with the literature[31], we identified the organic component was triglyceride (Table S6, Fig. S8). Osteoporosis has been associated with higher fat contents, especially in the bone marrows. For example, an inverse relationship between bone marrow adiposity and bone mass has been discovered in OP[32]. However, the marrows and any associated soft tissues on bone were removed carefully from our SSNMR sample, and any fat diffused into bone minerals from marrow would only be minor. We conclude that triglyceride is the special mobile molecule that is closely associated with OP induced by aging, especially in porous cancellous lumbar bone. Furthermore, our freeze-dried sample preparation also suggest the mobility of triglyceride is not from the suspension of triglyceride in bulk water since the bulk water is removed.

**Figure 10.**
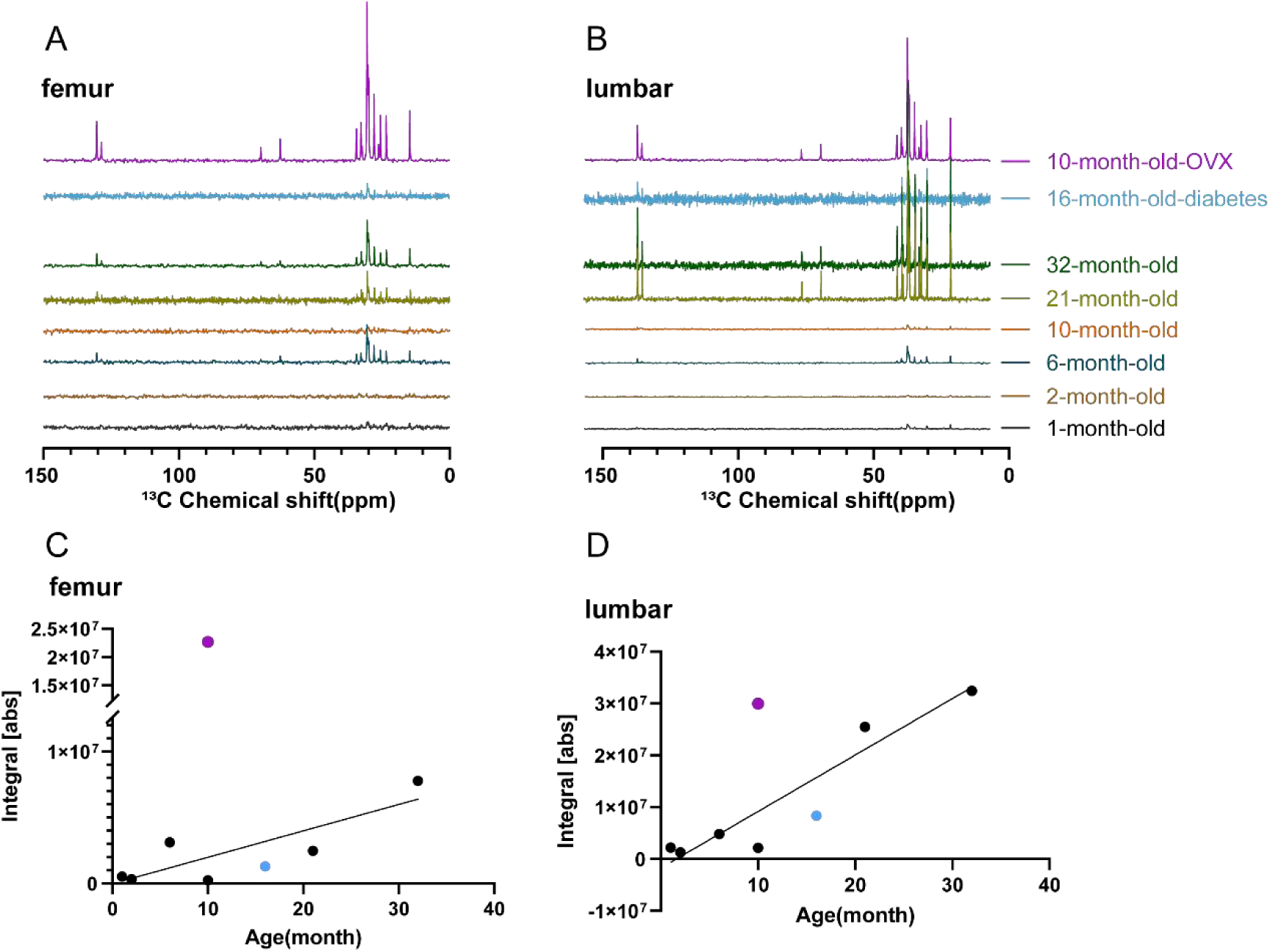
1D ^1^H-^13^C INEPT spectral changes with age and diseases. 1D ^1^H-^13^C INEPT spectra for (A) femur and (B) lumbar. The peak integral from 10 ppm to 40 ppm, relating to the amounts of triglyceride in samples, were shown in (C) femur; (D) lumbar. All the analysis were integral per unit mass. Purple: the 10-month-old-OVX SD rat; Blue: the 16-month-old male diabetic SD rat.

### Ovariectomy-induced osteoporosis SD rat model and type 2 diabetic SD rat model

In order to gain a better understanding of osteoporosis from different causes, we selected a 10-month-old female SD rat that underwent ovariectomy (OVX) at 6 months of age to simulate postmenopausal primary osteoporosis caused by estrogen deficiency. Another model was a 16-month-old male SD rat with induced type 2 diabetes since it is a disease frequently accompanied by a high risk for osteoporosis. This is also only male SD rat studied in this research. *μ*-CT results show a significant increase in the trabecular space only for cancellous bone in both the OVX and diabetic model (Fig. 2, 3 and Fig S1) comparing to bones with similar ages. Quantitative analysis of cancellous bone in OVX and diabetic model demonstrate a decrease in BV/TV and BMD, with both values much lower than those expected in their ages (compared to rat samples at 10 months and 21 months) (Fig. 4C, D). The results were different for cortical femur bone, with both models exhibiting relative bigger values of BV/TV and BMD than those corresponding to their ages (Fig. 4A, B). For skull, OVX model displayed bigger values of BV/TV and BMD while diabetic model displayed reduced values of BV/TV and BMD compared to similar age rat bones (Fig. 4E, F). Therefore, lumbar bone displayed more evident and characteristic changes of osteoporosis for both models, which is similar to that caused by aging. The OVX and diabetic models could show changes indicating osteoporosis in lumbar and skull but not in femur.

The solid-state NMR 2D ^1^H-^31^P HETCOR spectra of femur, lumbar and skull samples for both models were shown in Fig. S7 and the quantitative analysis results showed in fig 6B, C, D. The results from the three types of bones were similar. For OVX model at 10 months, the contents of HAP and ACP are very low, with a value similar to that of 6 months’ old rats for cortical bone or 2 months’ old rats for skull, or even lower for the cancellous lumbar bone. Therefore, OVX caused OP exhibits a significant low content of rigid minerals HAP and ACP. SD rats at 6 months should be at a stage of maturing according to μ-CT BV/TV and BMD (Fig 4). However, our SSNMR results do indicate ACP reaching maximum at 10 months while HAP at 21 months. Therefore, even the bone volume or bone mineral density reaches the maximum, the chemical contents would continue to develop to an older age. These extreme low values of HAP and ACP for OVX model are probably caused by the low starting values of HAP and ACP at 6 months when OVX is done, which deprives of the possibility of further growth in bones.

For the diabetic model at 16 months, HAP content is close to a value corresponding to the sample’s age for cortical femur and skull. However, ACP values decrease significantly compared to the normal aged curve for all three types of bones. Therefore, it is indicated osteoporosis associated with type 2 diabetes exhibits a sever decrease in ACP content, but not in HAP content compared to normal aging. The 1D ^31^P CP linewidth for both disease models displays similar or lower values than the expected values at the corresponding ages (Fig. 7A, C, E and Fig. S4). And the proportion of HAP in total HAP and ACP intensity all show higher values than the expected values at the corresponding ages (Fig. 7B, D and F). These results from molecular level by NMR are consistent with an increase in the crystallinity in the minerals for both animal models, especially for diabetes, which may cause bone brittle and easy to break.

As to the organic type I collagen in the bones,1D ^1^H-^13^C CP spectroscopy results for the OVX model show no significant difference in its content in cortical, cancellous and skull bones compared to normal aging groups (Fig. 8C, D; Fig. S5B). However, the 1D ^1^H-^13^C CP spectrum for the diabetic model lumbar bone shows a difference in the relative peak intensities around 20-50 ppm region, indicating a change in the organic components. 1D ^1^H DP experiments and 1D ^1^H-^13^C INEPT display the content of the mobile organic component triglyceride on cortical and cancellous bone for OVX and diabetic samples. The results consistently show an increase in its content for all three types of bones for OVX model comparing to bones at the same age (Fig. 9C, D; Fig. 10C, D; Fig. S5D, F). The increase is more significant for femur and skull, with the values bigger than those obtained for 21-month-old, 32-month-old rat bones. For lumbar, the signal intensity is only slightly higher than the 10-month-old rat, but the absolute intensity is close to the integral value of femur OVX model. Although the skull bone does not show any increase in triglyceride content during normal aging, it displays such an increase for OVX model.The diabetic model shows an overall low intensity for the mobile organic component. Only lumbar displays a slight increase in triglyceride content, similar to the change caused by aging. Therefore, it is concluded that triglyceride in bone tissue as a mobile component associated with aging is more significant in OVX caused OP.

## Discussion

In this research, bone tissue changes upon aging and for two disease models were thoroughly characterized by combining of μ-CT and SSNMR. *μ*-CT displays bone volume and density changes while SSNMR displays information at the molecular level on bone minerals and organic components. Our data analysis uniquely combined BMD mean from *μ*-CT and integral of peaks of SSNMR and gave a quantitative description of mineral component changes during OP development. It can be concluded that cancellous bones suffer a severe loss in bone volume and density, causing fragile bones upon aging or diseases. However, the cortical bones show a continuously increase of HAP until 21 months of age, and the crystallinity increase is probably the cause of brittle bones upon aging. In the diabetic disease model, the increase of HAP is in combination with a consistent decrease of ACP in all three types of bones, which would cause a relatively bigger increase of HAP and brittleness of bone than normal aging.

For OVX model, the contents for both HAP and ACP are very low, significant different from normal aging rats. We propose that although it is normally accepted that rats at 6 months of age reach their developing mature stage[33], also suggested by our *μ*-CT on cortical bone, the bone is still growing and developing which is clearly indicated by *μ*-CT on cancellous bone and SSNMR studies on the individual mineral molecules HAP and ACP. Therefore, a slight postpone of OVX operation time to a later month (10 month) could be a choice in the future experiments.

On the other hand, the triglyceride increases significantly for all three types of bones for OVX rats, much higher than normal aging rats. However, the amounts of triglyceride in diabetic rats are similar to or lower than normal rats at the similar age. It is also interesting to notice that the mobile triglyceride increases are not observed for skull upon aging, but clearly shown for femur and lumbar, and only for OVX rats the increases are shown for all three types of bones. These observations again support that the triglyceride is closely related to OP and only certain diseases, not for all bones studies, excluding the possibility that it is merely from marrow adipose diffused into the bone. Evidence implicate adipocyte metabolism and fat topography are altered in type 2 diabetes [15, 34–36]. The diabetic rat also had a high-fat diet. Interestingly, we did not observe bigger changes of triglyceride in diabetic rat bone tissues. The diabetic rat is also the only male rat studied. The different sex could also be a reason to explain the different changes in these parameters [37].

Triglyceride or fatty acid have been associated with OP in different researches [14, 32, 38], but a few work has analyzed it in hard mineralized bone tissues[31, 32, 39, 40]. The mineralized tissue contains lipids at much lower levels than marrow. They can be loosely-bound/easy extractable or tightly-associated lipids. Loosely-bound lipids could be from bone cells (mostly osteocytes representing 90-95% of total bone cells, and triglyceride being a senescence maker of osteocyte) and blood circulation irrigating the mineralized tissue, while tightly-associated lipids are those complexed with macromolecules and minerals in the mineralized extracellular matrix[40]. How thoroughly the marrow is removed from the bone and how thoroughly the lipids are extracted from the bone are two major issues in the past studies[41]. In our SSNMR study, the lipid extraction step is not needed, avoiding a potential error source. Using ^1^H-based MAS SSNMR, Mroue et. al has detected and characterized the mobile triglyceride in compact bovine cortical bone, indicating that it is from lipid embedded on the collagen fibrils [31]. Previous AFM (atomic-force microscopy) study on cortical bone of bovine tibia showed a layer of round particles covering the collagen bundles after demineralization. The particles could be dissolved in chloroform-ethanol solution, suggesting it is lipids. The lipids extracted from bones show a high percentage of triglycerides and cholesterol esters, which was correlated to the lipoproteins identified in bones using different compositional analysis techniques in that research [42]. Lipids could also be signal molecules, regulation the remodeling of bone, or involved in other processes. Although the exact function of triglyceride is beyond our current research, our results clearly indicate the increase of triglyceride in mineralized bone tissue is correlated to OP, especially OVX induced OP at young rat age.

In summary, our research gives a new and comprehensive picture on how osteoporosis with different causes is different, providing important parameters and molecular marks in diagnosis. The research also provides directions of future investigation to understand the mechanism of changes of these parameters.

## Acknowledgements

The work is supported by grants from the Natural Science Foundation of China (No. 32171185, 31770790 to J.-X.L.), Natural Science Foundation of Shanghai (20ZR1437000 to Y.Z.) and startup funding from ShanghaiTech University (2019F0202-000-01 to Y.Z.). We would like to thank the Biomolecular NMR Facility at the School of Life Science and Technology, ShanghaiTech University for SSNMR studies.

## Reference

1. Zimmermann, E.A., et al., Age-related changes in the plasticity and toughness of human cortical bone at multiple length scales. Proceedings of the National Academy of Sciences of the United States of America, 2011. 108(38): p. 14416.

2. Lane, J.M., L. Russell, and S.N. Khan, Osteoporosis. Clin Orthop Relat Res, 2000(372): p. 139–50.

3. Clarke, B., Normal bone anatomy and physiology. Clin J Am Soc Nephrol, 2008. 3 **Suppl 3**(Suppl 3): p. S131-9.

4. Bouxsein, M.L., et al., Guidelines for assessment of bone microstructure in rodents using micro-computed tomography. J Bone Miner Res, 2010. 25(7): p. 1468–86.

5. Campbell, G.M. and A. Sophocleous, Quantitative analysis of bone and soft tissue by micro-computed tomography: applications to ex vivo and in vivo studies. Bonekey Rep, 2014. 3: p. 564.

6. Bolamperti, S., I. Villa, and A. Rubinacci, Bone remodeling: an operational process ensuring survival and bone mechanical competence. Bone Res, 2022. 10(1): p. 48.

7. Dorozhkin, S.V. and M. Epple, Biological and medical significance of calcium phosphates. Angew Chem Int Ed Engl, 2002. 41(17): p. 3130–46.

8. Lowenstam, H.A. and S. Weiner, On Biomineralization. 1989: Oxford University Press.

9. Viguet-Carrin, S., P. Garnero, and P.D. Delmas, The role of collagen in bone strength. Osteoporos Int, 2006. 17(3): p. 319–36.

10. Reznikov, N., R. Shahar, and S. Weiner, Bone hierarchical structure in three dimensions. Acta Biomater, 2014. 10(9): p. 3815–26.

11. Zeng, P., et al., Solid-State Nuclear Magnetic Resonance Identifies Abnormal Calcium Phosphate Formation in Diseased Bones. ACS Biomater Sci Eng, 2021. 7(3): p. 1159–1168.

12. Mroue, K.H., et al., Proton-Detected Solid-State NMR Spectroscopy of Bone with Ultrafast Magic Angle Spinning. Sci Rep, 2015. 5: p. 11991.

13. Cheng, C.H., L.R. Chen, and K.H. Chen, Osteoporosis Due to Hormone Imbalance: An Overview of the Effects of Estrogen Deficiency and Glucocorticoid Overuse on Bone Turnover. Int J Mol Sci, 2022. 23(3).

14. Zhang, J., et al., The role of lipid metabolism in osteoporosis: Clinical implication and cellular mechanism. Genes Dis, 2024. 11(4): p. 101122.

15. Chau, D.L. and S.V. Edelman, Osteoporosis and Diabetes. Clinical Diabetes, 2002. 20(3): p. 153–157.

16. Yousefzadeh, N., et al., Ovariectomized rat model of osteoporosis: a practical guide. Excli j, 2020. 19: p. 89–107.

17. Jiao, H., E. Xiao, and D.T. Graves, Diabetes and Its Effect on Bone and Fracture Healing. Curr Osteoporos Rep, 2015. 13(5): p. 327–35.

18. Abdulameer, S.A., et al., Osteoporosis and type 2 diabetes mellitus: what do we know, and what we can do? Patient Prefer Adherence, 2012. 6: p. 435–48.

19. Yang, C., et al., 3D Printed Enzyme 〧unctionalized Scaffold Facilitates Diabetic Bone Regeneration. Advanced Functional Materials, 2021.

20. Morcombe, C.R. and K.W. Zilm, Chemical shift referencing in MAS solid state NMR. Journal of Magnetic Resonance, 2003. 162(2): p. 479–486.

21. Fung, B.M., A.K. Khitrin, and K. Ermolaev, An Improved Broadband Decoupling Sequence for Liquid Crystals and Solids. Journal of Magnetic Resonance, 2000. 142(1): p. 97–101.

22. Lesage, A. and L. Emsley, Through-Bond Heteronuclear Single-Quantum Correlation Spectroscopy in Solid-State NMR, and Comparison to Other Through-Bond and Through-Space Experiments. Journal of Magnetic Resonance, 2001. 148(2): p. 449–454.

23. Kocbach, L. and S. Lubbad, Geometrical simplification of the dipole-dipole interaction formula. Physics Education, 2010. 45(4): p. 345–345.

24. Morris, G.A. and R. Freeman, Enhancement of nuclear magnetic resonance signals by polarization transfer. Journal of the American Chemical Society, 1979. 101(3): p. 760–762.

25. Shaka, A.J., et al., An improved sequence for broadband decoupling: WALTZ-16. 1969. 52(2): p. 335–338.

26. Andrew, E.R., Nuclear Magnetic Resonance Spectra from a Crystal rotated at High Speed. Nature, 1958. 182(4650): p. 1659.

27. Vega and J. Alexander, Heteronuclear chemical-shift correlations of silanol groups studied by two-dimensional cross-polarization magic angle spinning NMR. Journal of the American Chemical Society, 1988. 110(4): p. 1049–1054.

28. Cao, A.B., L.M. McGrady, and M. Wang, *Effect of age on femur whole-bone bending strength of mature rat.* Clin Biomech (Bristol, Avon), 2023. 101: p. 105828.

29. Wang, H., et al., Biomechanical properties and clinical significance of cancellous bone in proximal femur: A review. Injury, 2023.

30. Emmanuelle, N.E., et al., Critical Role of Estrogens on Bone Homeostasis in Both Male and Female: From Physiology to Medical Implications. Int J Mol Sci, 2021. 22(4).

31. Mroue, K.H., et al., Selective detection and complete identification of triglycerides in cortical bone by high-resolution (1)H MAS NMR spectroscopy. Phys Chem Chem Phys, 2016. 18(28): p. 18687–91.

32. During, A., Osteoporosis: A role for lipids. Biochimie, 2020. 178: p. 49–55.

33. Quinn, R., Comparing rat’s to human’s age: how old is my rat in people years? Nutrition, 2005. 21(6): p. 775–7.

34. Bays, H., L. Mandarino, and R.A. DeFronzo, Role of the adipocyte, free fatty acids, and ectopic fat in pathogenesis of type 2 diabetes mellitus: peroxisomal proliferator-activated receptor agonists provide a rational therapeutic approach. J Clin Endocrinol Metab, 2004. 89(2): p. 463–78.

35. Picke, A.K., et al., Update on the impact of type 2 diabetes mellitus on bone metabolism and material properties. Endocr Connect, 2019. 8(3): p. R55–r70.

36. Blank, R.D., et al., Association between type 2 diabetes and osteoporosis risk: A representative cohort study in Taiwan. Plos One, 2021. 16(7).

37. Zhang, Y.Y., et al., Insights and implications of sexual dimorphism in osteoporosis. Bone Res, 2024. 12(1): p. 8.

38. Chang, P.Y., et al., Triglyceride Levels and Fracture Risk in Midlife Women: Study of Women’s Health Across the Nation (SWAN). J Clin Endocrinol Metab, 2016. 101(9): p. 3297–305.

39. Schulz, J., et al., Quantitative monitoring of extracellular matrix production in bone implants by 13C and 31P solid-state nuclear magnetic resonance spectroscopy. Calcif Tissue Int, 2007. 80(4): p. 275–85.

40. During, A., G. Penel, and P. Hardouin, Understanding the local actions of lipids in bone physiology. Prog Lipid Res, 2015. 59: p. 126–46.

41. During, A., Lipid determination in bone marrow and mineralized bone tissue: From sample preparation to improved high-performance thin-layer and liquid chromatographic approaches. J Chromatogr A, 2017. 1515: p. 232–244.

42. Xu, S. and J.J. Yu, Beneath the minerals, a layer of round lipid particles was identified to mediate collagen calcification in compact bone formation. Biophys J, 2006. 91(11): p. 4221–9.

