## Supplementary Figures 1-8 for "Osteoporosis with different causation exhibits different changes in bone tissue mineral and organic matrix"

Figure S1. The Micro-CT images of the skull, showing a progressive change of bone with age and OP. The sample was from 1, 2, 3, 6, 10, 21 and 32-month-old SD rat, 10-month-old-OVX and 16-month-old male SD rat with diabetes. The scale bar in each image is 500μm.

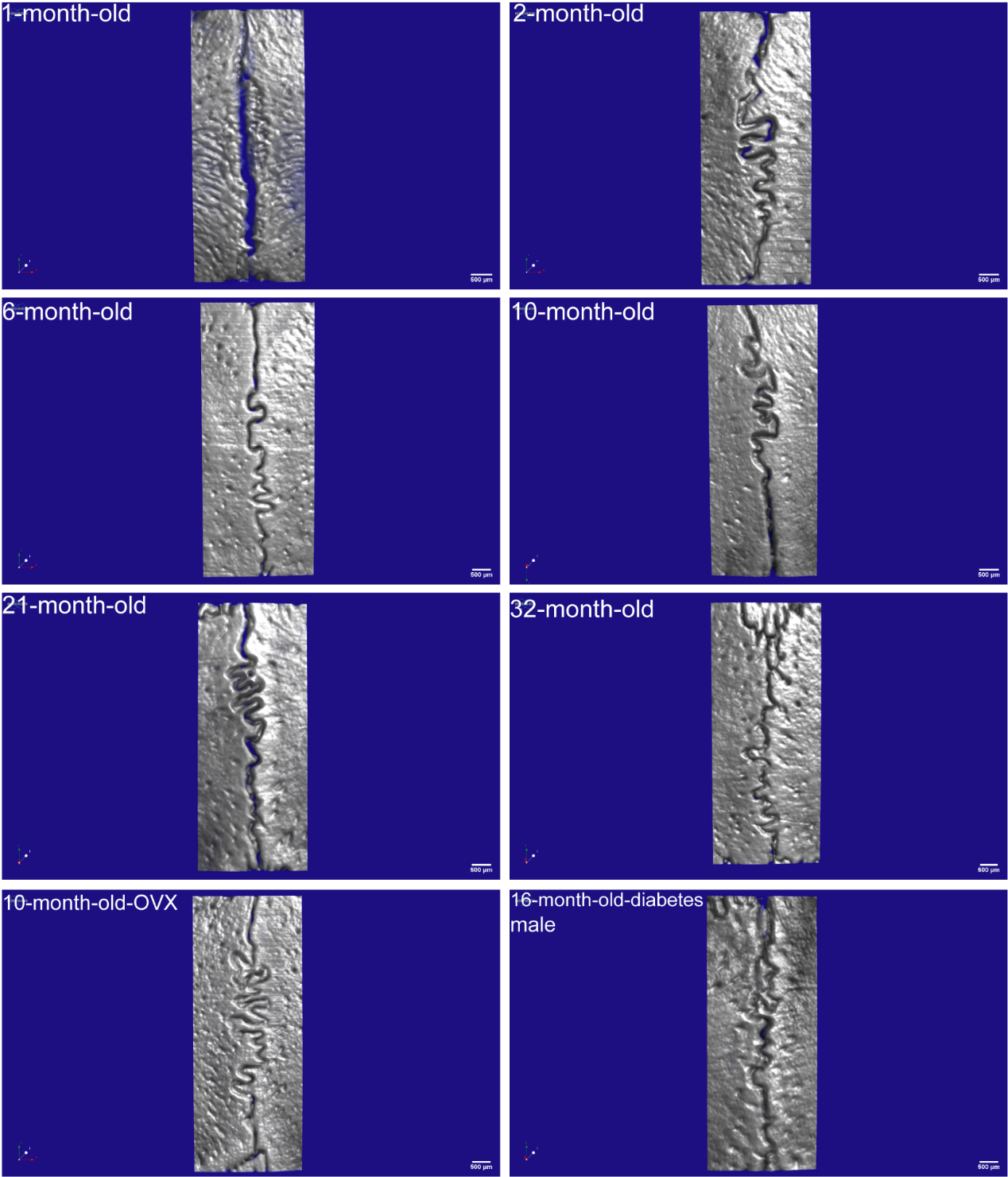

Figure S2. 2D ^1^H-^31^P HETCOR spectra of lumbar bones. A, B, C, D, E and F represent lumbar from 1, 2, 3, 6, 10, 21 and 32-month-old SD rat, respectively. All lumbar samples were freeze-dried, and the spectra were collected from 128 scans.

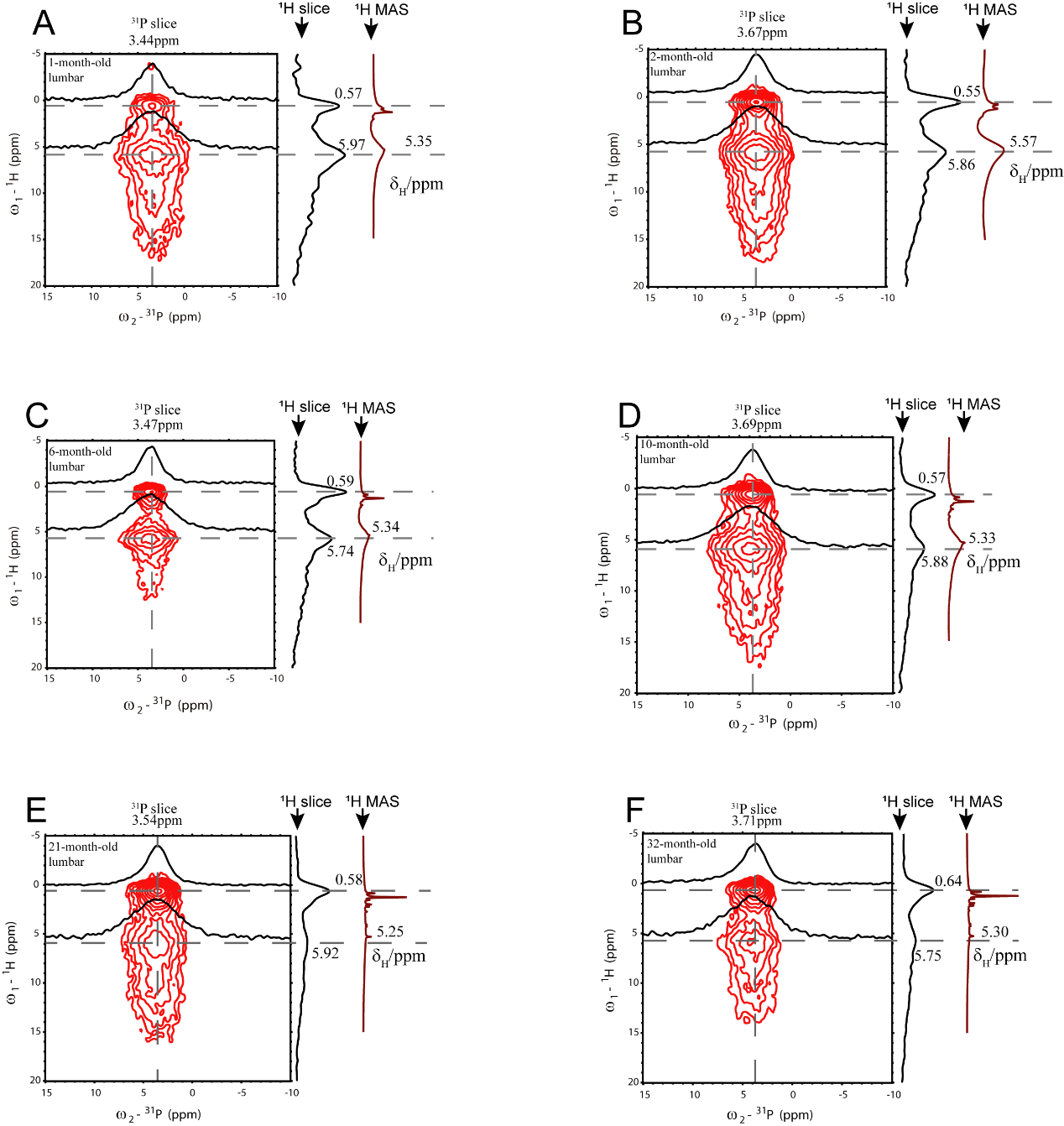

Figure S3. 2D ^1^H-^31^P HETCOR spectra of skull bones. A, B, C, D, E and F represent skull from 1, 2, 3, 6, 10, 21 and 32-month-old SD rat, respectively. All samples were freeze-dried, and the spectra were collected from 128 scans.

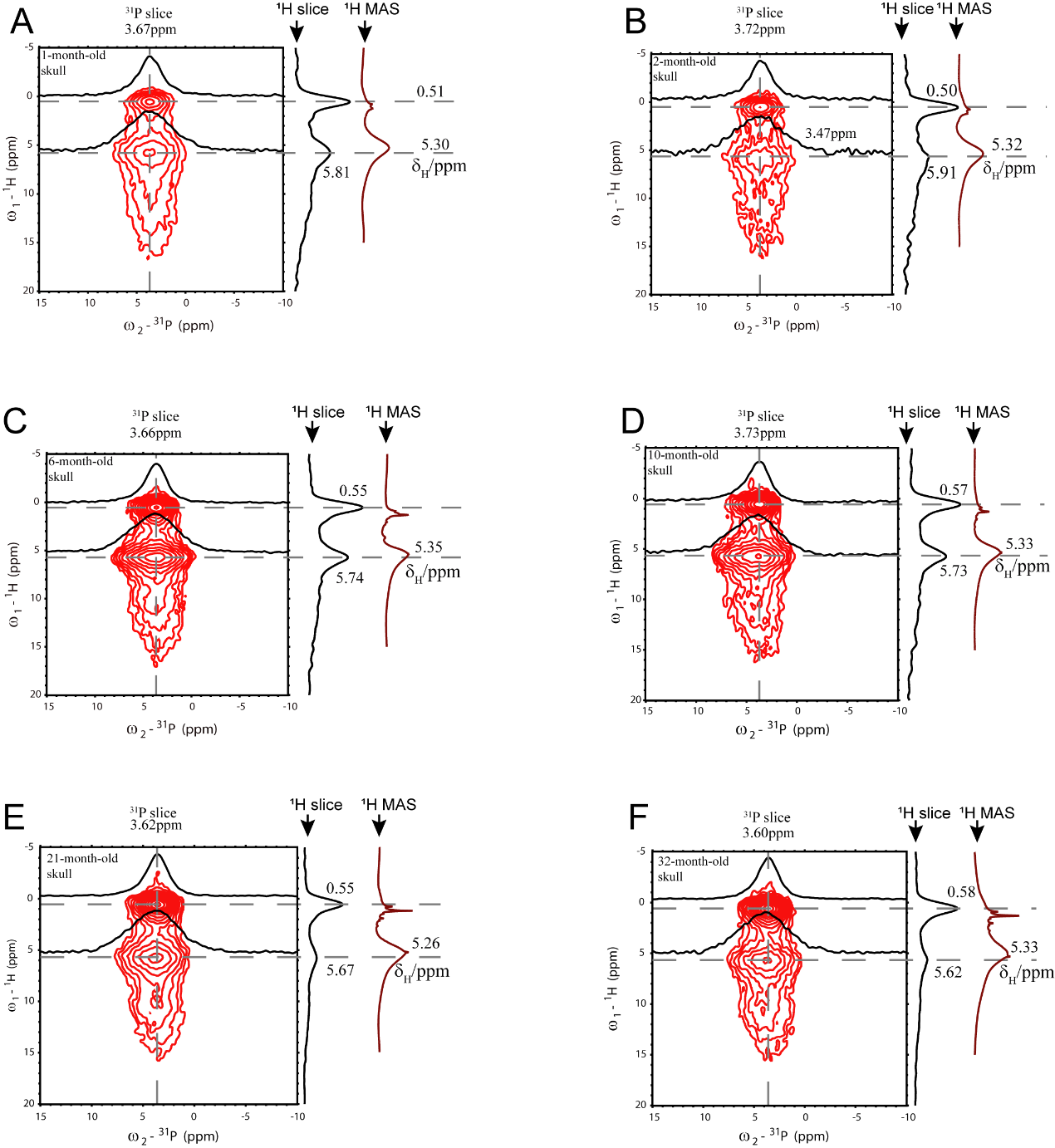

Figure S4. 1D ^31^P CP spectra of femur samples at 15kHz depicts a gradual decrease of the peak width at half height with aging and OP development. The exact values were shown in Table S1 for all the samples. The intensity of all signals in this figure were adjusted to the same height.

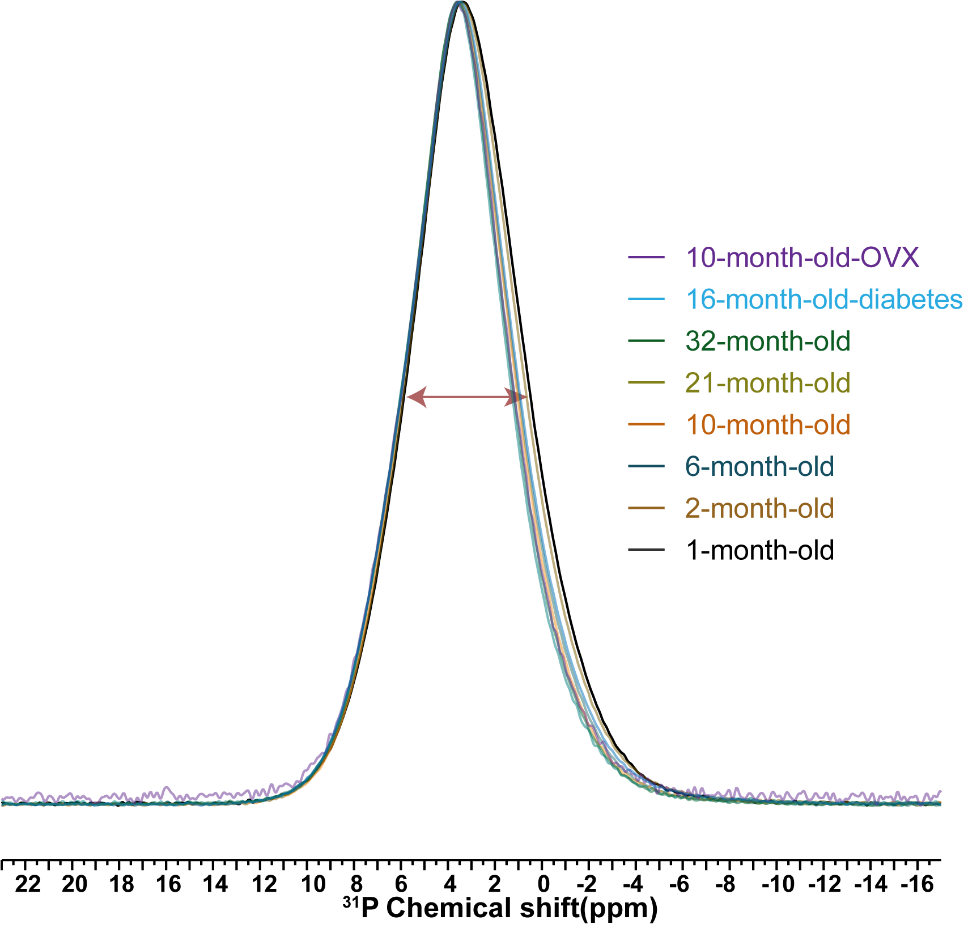

Figure S5. Aged-related changes of organic matter in the skull. A. 1D ^1^H-^13^C CP spectra of skull samples with different ages. B. The whole spectra integral changes as a function of rat age, reflecting the amount of collagen changes. All the analysis was integral per unit mass from 0 ppm to 200 ppm. C. 1D ^1^H DP spectra of skull samples with different ages and D. 1D ^1^H DP spectra integral changes as a function of rat age. All the analysis was integral per unit mass from 0 ppm to 3 ppm. E. 1D INEPT spectra of skull samples show the signal changes with age. F. 1D INEPT SSNMR spectral intergrals as a function of age. All the analysis was integral per unit mass from 10 ppm to 40 ppm. The results on disease models were also shown with results on the 10-month-old-OVX rat in purple and the16-month-old male diabetic SD rat in blue. All experiments were measured at 15kHz MAS.

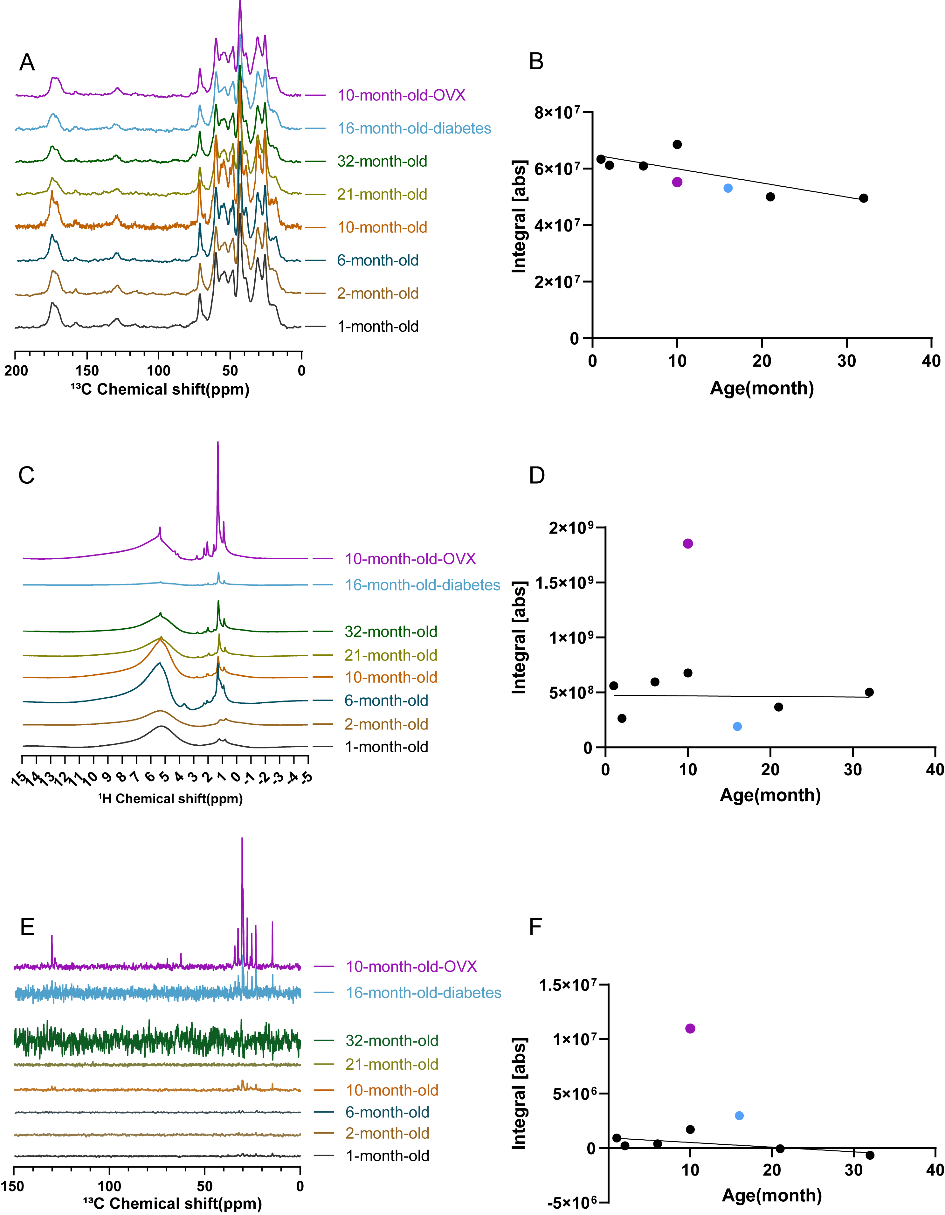

Figure S6. The ^1^H T_2_ values were obtained by exponentially fitting of the intensity of ^1^H peaks versus CPMG echo time. The values for various triglyceride peaks in a dehydrated 32-month-old lumbar sample were presented, which were also summarized in Table S2 with assignments.

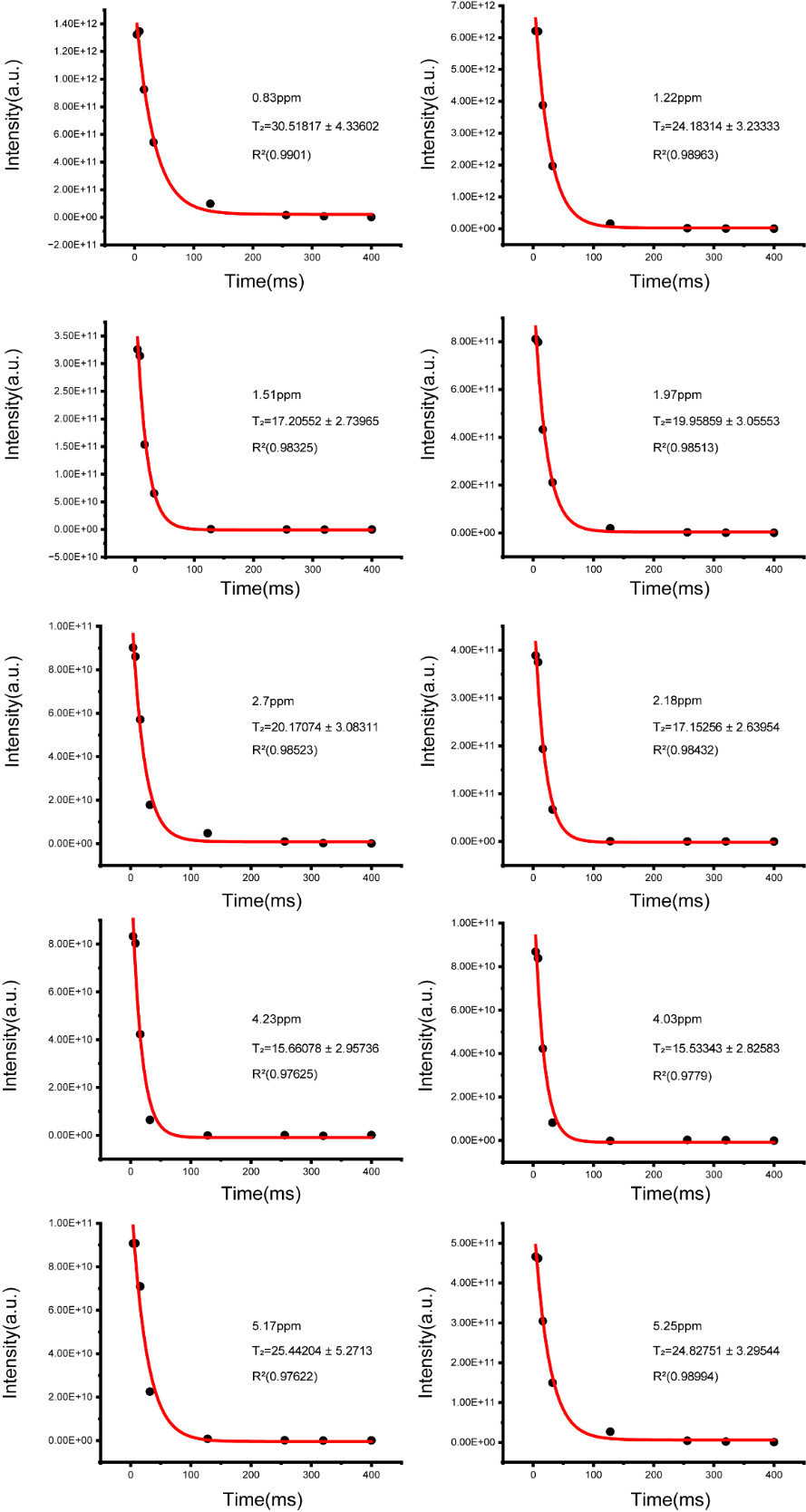

Figure S7. ^1^H-^31^P 2D HETCOR spectra of bones from disease models. A, C and E were from femur, lumbar and skull samples of a 10-month-old-OVX SD rat. B, D and F were from femur lumbar and skull samples of a 16-month-old male diabetic SD rat. The bone samples were freeze-dried and the number of scans were 128 for each spectrum.

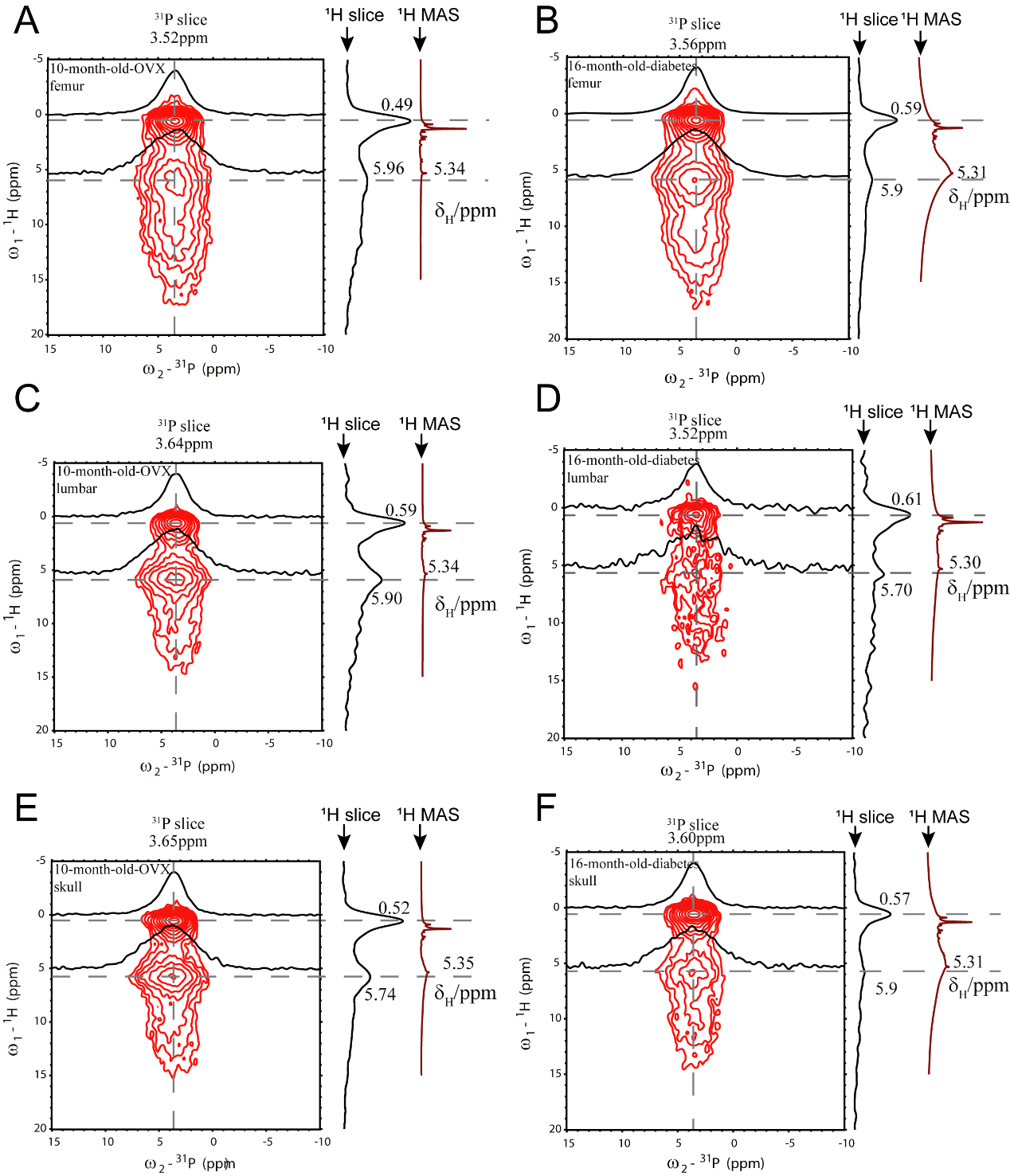

Figure S8. 2D ^1^H-^13^C INEPT spectra of three bone samples from a 10-month-old-OVX rat. (A) The chemical structure of triglycerides with the carbon atom numbering matching to those shown in the spectra for (B) lumbar. 2D ^1^H-^13^C INEPT spectrum of (C) femur and (D) skull were similar to (B).

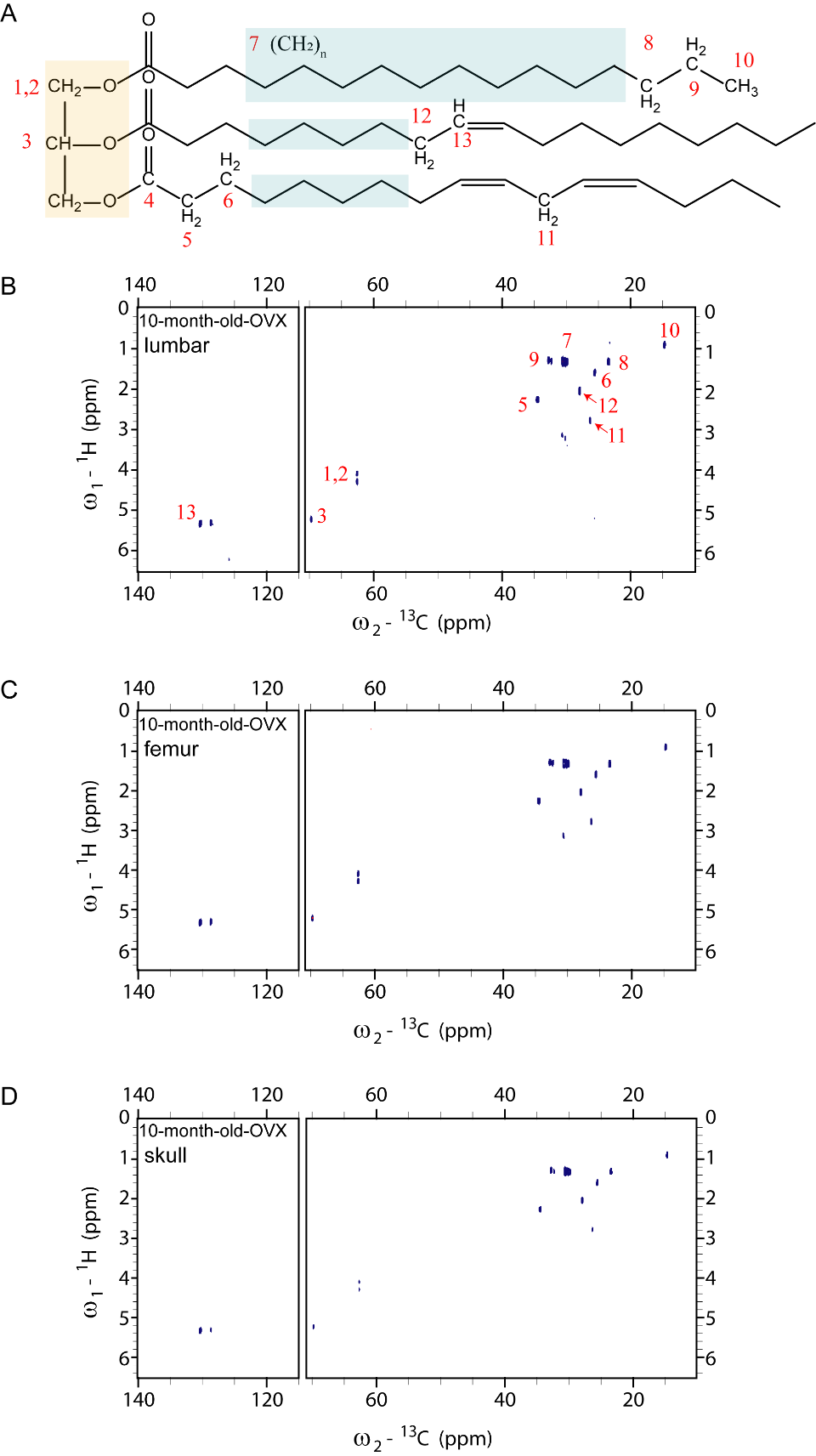
